# Diacylglycerol kinase ζ dictates CD40-mediated immune synapse formation, mTORC1 signaling and plasma cell fate in B lymphocytes

**DOI:** 10.1101/2025.01.11.632541

**Authors:** Ana Fernández-Barrecheguren, Adrián Fernández-Rego, Lucía Fuentes-Cantos, Tirso Pons, Belén S. Estrada, Marta Iborra-Pernichi, Tahmineh Ebrahimi, Ana Sagrega-Aparisi, Sara Cogliati, Nuria Martínez-Martín, Rodrigo Jiménez-Saiz, Yolanda R. Carrasco

## Abstract

To mount a robust T-dependent immune response, antigen-specific B lymphocytes require CD40 stimulation through immune synapse formation with CD4^+^ T follicular helper cells. CD40 triggers the activation of mammalian target of rapamycin complex-1 (mTORC1) and remodels the mitochondria to meet increased bioenergetic and anabolic demands. We show that diacylglycerol-kinase-ζ (DGKζ) has a crucial role in activating the mTORC1 pathway and remodeling mitochondria downstream of CD40 signaling in B cells. DGKζ governs organelle translocation to the CD40-mediated immune synapse and the recruitment of mTORC1 to lysosomes. DGKζ^−/−^ B cells exhibited impaired mitochondria function, protein biosynthesis, metabolite transporter expression and cell cycle progression, accompanied by dysregulation of the transcriptional network governing B cell fate. These defects lead to a blockage in the progression of the germinal center response and plasma cell differentiation *in vivo*. Our findings establish DGKζ as a key mediator of CD40 functions in the B cell response.

## INTRODUCTION

Upon recognizing an antigen *via* the B cell antigen receptor (BCR), B cells become activated within secondary lymphoid organ follicles. This involves the formation of an immune synapse with an antigen-presenting cell, enabling stable cell-cell interactions, signaling, organelle polarization and antigen uptake^1–4^. Activated B cells subsequently migrate to the follicular border, where they present processed antigen to Tfh cells on MHC class II molecules. In return, Tfh cells provide co-stimulatory signals, primarily through CD40 stimulation, driving B cell survival and proliferation^5^. While the molecular mechanisms of this immune synapse have been extensively studied from the T cell perspective, its structure and function on the B cell side remain largely unknown. The CD40/CD40 ligand (CD40L) axis is a key checkpoint for a robust humoral immune response against pathogens. CD40, a member of the TNF receptor superfamily, is constitutively expressed as a homodimer on the B cell surface. Upon binding of trimeric CD40L, CD40 trimerizes, initiating a cascade of downstream signaling events^6–8^. Fully activated B cells can differentiate into short-lived antibody-secreting plasma cells and memory B cells (MBCs). Alternatively, they can migrate to germinal centers to generate high-affinity, long-lived plasma cells and MBCs. CD40 signaling promotes immunoglobulin class switching and somatic hypermutation processes, essential for BCR affinity maturation^9–11^.

Activated B cells undergo a metabolic shift to meet the increased energy and biosynthetic demands required for cell growth, proliferation and differentiation. This is primarily driven by the activation of the mTORC1 pathway in conjunction with enhanced mitochondrial respiration (oxidative phosphorylation, OxPhos) and glycolysis^12–15^. Mitochondrial remodeling provides activated B cells with ATP and essential intermediate metabolites, such as those generated through the tricarboxylic acid cycle, to sustain their bioenergetic and anabolic needs^16–18^. mTORC1, a serine-threonine kinase, senses nutrient availability and regulates cell growth and division. It is activated by the PI3K/Akt pathway and nutrient signals, which facilitate the recruitment of mTORC1 to the lysosomal surface *via* RHEB and Rag GTPases, respectively, promoting biosynthetic processes^19,20^. These nutrient signals can include amino acids, glucose and phosphatidic acid (PA), a crucial metabolite for phospholipid biosynthesis. PA is essential for mTORC1 stability and function^21^. While various sources can provide PA, DGKζ-derived PA has been shown to promote mTORC1 activation in cancer cells and other cell types^22,23^. However, the role of DGKζ in mTORC1 activation and metabolic reprograming of activated B cells remains largely unexplored.

DGKζ is a member of the DGK family of enzymes, which phosphorylate the lipid messenger diacylglycerol (DAG) to generate PA. As one of the most extensively studied DGK isoforms, DGKζ possesses a complex domain structure that allows DGKζ to function both as a kinase and an adapter protein. Phosphorylation by PKC triggers the translocation of DGKζ from the cytosol to membrane interfaces where DAG is produced. The C1 domains further facilitate the targeted localization of DGKζ to specific membranes through DAG binding^24–26^. DGKζ is known to suppress the function of lymphocytes by restricting signaling pathways activated by antigen receptor stimulation, specifically those dependent on DAG. Furthermore, the local production of PA serves as a nucleation site to recruit effector proteins involved in cytoskeletal remodeling and cell polarity^26,27^. Actomyosin remodeling, integrin clustering and polarized membrane trafficking are key characteristics of the immune synapse, and we previously showed that all these processes are promoted by DGKζ at the BCR-mediated synapse, facilitating the efficient extraction of antigen^28^. The role of DGKζ in the co-stimulatory step *via* CD40, which requires the assembly of a synapse with Tfh cells and is indispensable for the germinal center response, remains unknown.

We therefore investigated the role of DGKζ in CD40 signaling. Using DGKζ-deficient mouse B cells and GFP-tagged DGKζ mutants, we found that DGKζ promotes mTORC1 activation following CD40 stimulation. This activation relies on both PA production and an adaptor function of DGKζ through its PDZ binding motif (PDZbm). CD40-stimulated DGKζ^−/−^ B cells generated fewer plasma cells than equivalent wild-type (WT) cells, correlating with dysregulated expression of transcription factors involved in plasma cell differentiation. Anabolism, cell cycle progression and mitochondrial function were also impaired in DGKζ^−/−^ B cells downstream of CD40 signaling. *In vivo* immunization experiments revealed a blockade in germinal center progression in DGKζ^−/−^ B cells. We dissected the CD40/DGKζ crosstalk at the B cell synapse with Tfh cells by recreating this interaction *in vitro*, finding that DGKζ promotes actin polymerization and LFA-1-mediated adhesion at the CD40-triggered synapse-like contact, while limiting B cell exploratory behavior. DGKζ also regulates organelle translocation to the synapse, facilitating mTORC1 recruitment to lysosomes and its subsequent activation. Our data thus establish DGKζ as a crucial mediator of the CD40 checkpoint in the B cell immune response.

## RESULTS

### DGKζ promotes mTORC1 activity and plasma cell generation downstream of CD40 signaling

To explore the role of DGKζ in the signaling of CD40 in B cells, we isolated B cells from the spleen of WT and DGKζ^−/−^ mice and stimulated them *in vitro* with an antibody against CD40. Both DGKζ^−/−^ and WT B cells expressed comparable levels of surface CD40, as determined by flow cytometry (Figure S1A). Analysis of the early signaling cascade by immunoblotting revealed the activation of PKCβ, Akt, and S6K1(p70-S6K) in WT B cells, along with phosphorylation of FOXO1 at Thr24, indicating the inactivation of transcriptional repressor functions. DGKζ^−/−^ B cells displayed significantly elevated levels of the activated forms of PKCβ and Akt when compared with WT cells, accompanied by higher levels of phosphorylated FOXO1 (Figures 1A-B). To evaluate mTORC1 activity, we monitored the phosphorylated status of ribosomal protein S6 (p-S6), a downstream target, by flow cytometry. Both WT and DGKζ^−/−^ B cells exhibited increased p-S6 levels upon CD40 stimulation, peaking at 30 min; however, p-S6 levels were significantly lower in DGKζ^−/−^ cells than in WT cells (Figure 1C). To reconstitute our system, we added soluble PA and employed the inhibitor rapamycin to monitor mTORC1 activity. We observed elevated levels of p-S6 in PA-treated DGKζ^−/−^ B cells at the 15-min time point (Figure 1D). This suggests that soluble PA can restore mTORC1 activity downstream of CD40 signaling in DGKζ^−/−^ B cells during the early phase of activation. To further investigate the relevance of the kinase *versus* the adaptor activities of DGKζ in mTORC1 regulation, we used the mouse A20 B cell line to express different DGKζ constructs (WT; kinase-dead, KD; and a deletion mutant for the PDZbm, ΔPDZbm) fused to GFP. A20 cells express CD40 on their surface and lack endogenous DGKζ (Figures S1B-C). We confirmed construct expression by GFP detection at 24 h (Figure S1D), and then analyzed p-S6 levels after CD40 stimulation (Figure S1E). The fraction of GFP-negative (GFP^−^) cells was analyzed as a control. Basal p-S6 levels increased in GFP^−^ cells at 15 min post-CD40 stimulation. While DGKζ-WT expression did not alter this pattern, both the DGKζ-KD and -ΔPDZbm mutants blunted the p-S6 response (Figure S1F). These findings indicate that both the kinase activity and the PDZbm-mediated adaptor function of DGKζ are involved in regulating mTORC1 activation *via* CD40 signaling.

**Figure 1.**
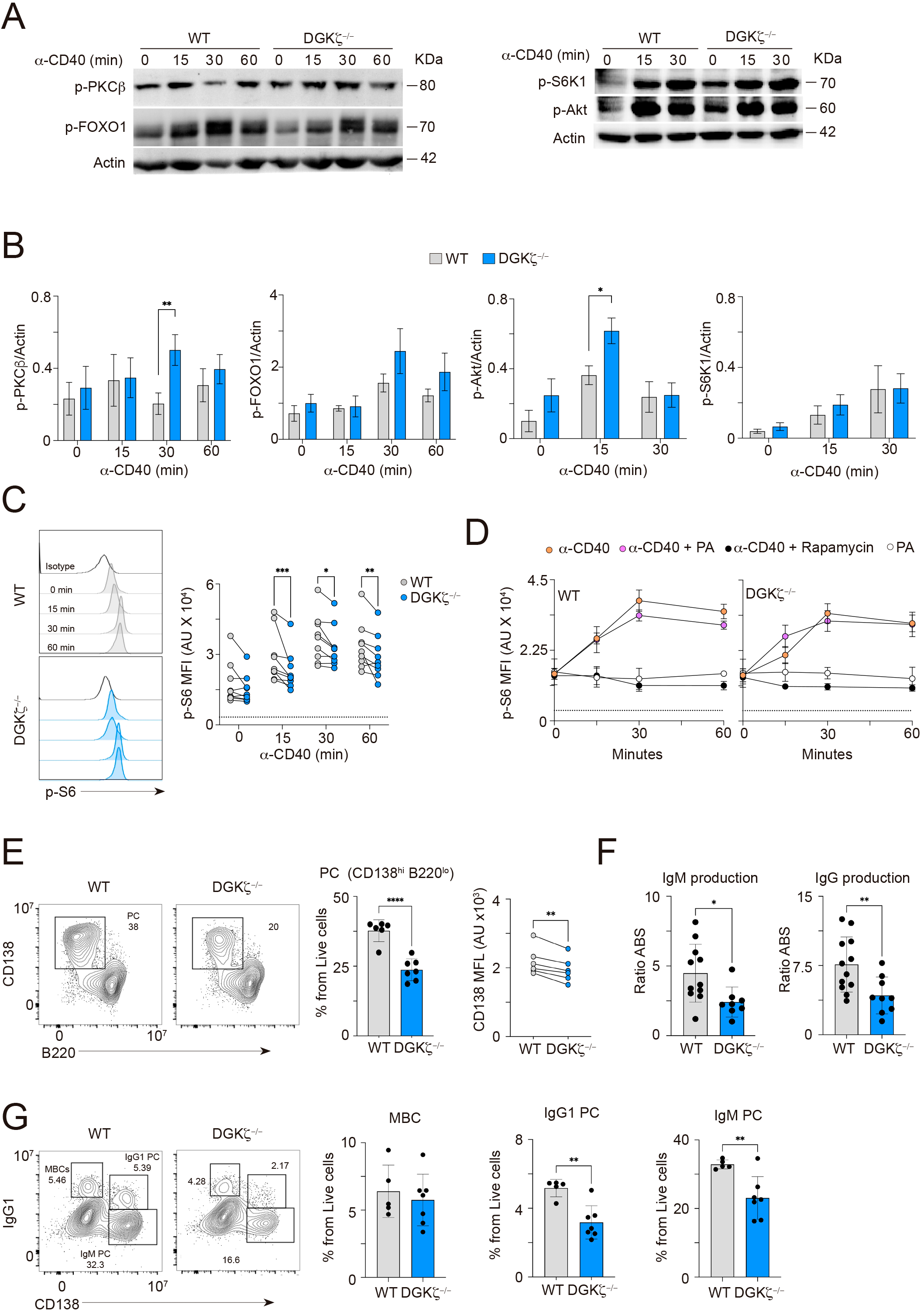
DGKζ promotes mTORC1 activity and plasma cell generation downstream CD40. **A-C,** B cells were stimulated with anti-CD40 antibody. **A**, Immunoblots for the phosphorylated forms (p-) of PKCβ (S660), FOXO1 (T24), S6K1 (T389) and Akt (S473); actin, loading control. **B**, Quantification of band intensities in A, normalized to actin. Data are the mean±SEM (n=3-5 mice). **C**, Profiles of the p-S6 (S235/236) and quantification of the mean fluorescence intensity (MFI) values in arbitrary units (AU) over time; each dot corresponds to one mouse. Dotted line, isotype control. **D**, as in C but adding phosphatidic acid (PA) or rapamycin to the assay. Data are the mean±SEM (n=3 mice). Dotted line, isotype control. **E-H**, B cells were cultured with anti-CD40/IL-4/IL-5 for 96 h and analyzed by flow cytometry (see Figure S1G); supernatants were used for IgM/IgG detection by ELISA. Each dot corresponds to one mouse. **E**, CD138/B220 contour-plots showing the plasma cell population (PC; CD138^hi^B220^lo^); percentages are indicated. Quantification of PC frequency and CD138 MFI values. **F,** Quantification of IgM and IgG production; absorbance (ABS) values were normalized to that of un-stimulated cells supernatants. **G,** IgG1/CD138 contour-plots showing IgG1^+^ MBCs, IgG1^+^ and IgM^+^ PC, and frequency values. Bar-graphs, mean±SEM. Statistical analysis: two-tailed unpaired (B, D, F) or paired (C, E, G, H) parametric Student’s t-test. *, *p*<0.05; **, *p*<0.01; ***, *p*<0.001; ****, *p*<0.0001.

To investigate the long-term effects of DGKζ on CD40 signaling, we cultured WT and DGKζ^−/−^ B cells for 96 h in the presence of anti-CD40 antibody, IL-4 and IL-5 and analyzed plasma cell generation. Results showed that the plasma cell population (CD138^+^B220^low/-^) was significantly lower in DGKζ^−/−^ cells than in WT controls, as was the surface expression of CD138, indicating impaired plasma cell maturation (Figures 1E and S1G). ELISA analysis revealed lower IgM and IgG production in the culture supernatant of DGKζ^−/−^ B cells (Figure 1F). No significant differences were observed in the IgG1^+^ MBC-like generation between the two genotypes, but both IgG1^+^ and IgM^+^ plasma cells were reduced in the DGKζ^−/−^ cells (Figure 1G). These findings suggest that DGKζ plays a crucial role in CD40-mediated B cell signaling by suppressing the PKCβ/Akt pathway, enhancing mTORC1 activity and promoting plasma cell differentiation.

### DGKζ is essential for the expression of transcription factors involved in plasma cell commitment

A complex network of transcription factors regulates plasma cell differentiation. PAX5, BCL6 and BACH2 are involved in B cell identity, whereas IRF4, BLIMP1 and XBP1 are involved in plasma cell commitment and function^29^. To investigate the impact of DGKζ on this regulatory network, we monitored mRNA expression levels of those genes in CD40-activated DGKζ^−/−^ and WT B cells, before the emergence of CD138^+^B220^low^ cells (Figure S2A). Quantitative RT-PCR analysis revealed that the expression of *Pax5* and *Bcl6* was higher in DGKζ^−/−^ B cells than in WT controls both at baseline and after 48 h of culture, while *Irf4* levels were significantly lower at 48 h (Figure 2A). To validate these findings, we performed RNA-seq analysis of the 48-h CD138^−^B220^+^CD95^+^GL7^+^ population (Figure S2B; see Supplemental Methods and Document S2), which confirmed the higher expression of B cell-identity genes (*Pax5, Bcl6*) and lower expression of plasma cell-related genes (*Irf4, Prdm1*) in DGKζ^−/−^ B cells (Figure 2B). While no significant differences were observed in the expression of *Foxo1*, *Bach2* or *c-myc* (Figures 2A-B), DGKζ^−/−^ B cells exhibited higher expression of *Aicda*, required for class switching and affinity maturation, and lower expression of other genes associated with DNA repair (*E2f1, Ezh2*) (Figure 2B). To gain insights into the dynamics of protein expression, we monitored PAX5, IRF4 and BLIMP1 expression by flow cytometry. Consistent with previous results^13^, WT B cells upregulated PAX5 and IRF4 expression following CD40 stimulation. By 72 h, an IRF4^high^PAX5^low^ population emerged, representing plasma cell-committed cells that also began to express BLIMP1 (Figures 2C-D). Quantification of the IRF4^high^PAX5^low^ and BLIMP1^+^PAX5^low^ populations at 96 h revealed significant impairment in the ability of DGKζ^−/−^ B cells to generate plasma cell-committed cells compared with WT controls (Figures 2C-D). Collectively, these findings suggest that DGKζ plays a critical role in regulating the transcriptional network that governs plasma cell fate downstream of CD40 signaling.

**Figure 2.**
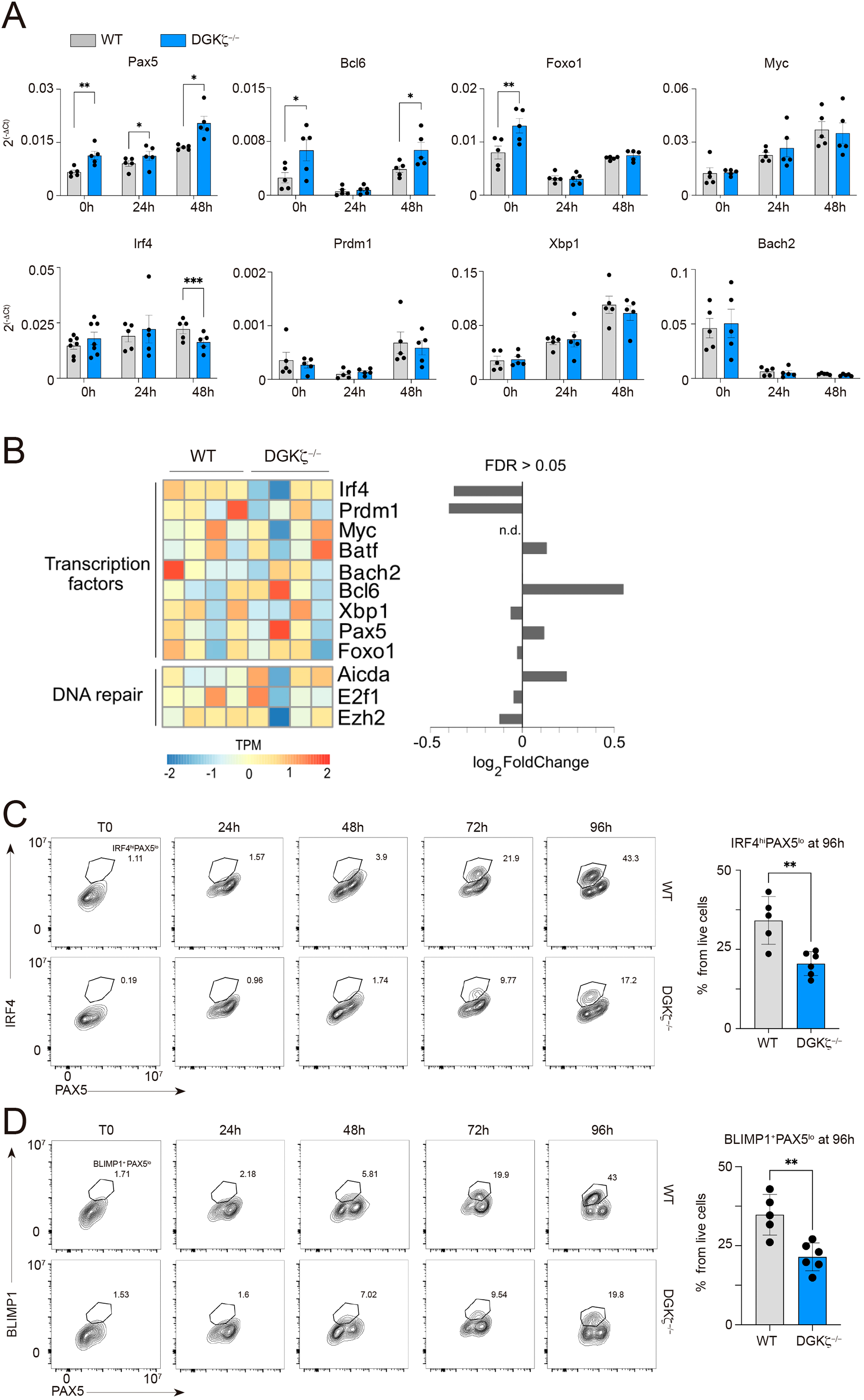
DGKζ is essential for the expression of transcription factors involved in plasma cell fate. **A,** B cells freshly isolated (0h) or cultured with anti-CD40/IL-4/IL-5 were analyzed for expression of indicated transcription factors by RT-qPCR; obtained values were normalized to actin expression. **B,** B cells cultured as in A for 48 h were analyzed by RNA-seq. Heatmap and bar-plot illustrating the expression profiles (TPM, transcript per million mapped reads) and differential gene expression (log_2_FoldChange) of selected transcription factors and DNA repair genes. Data from n=4 mice per genotype. FDR, the Benjamini-Hochberg False Discovery Rate; n.d., not determined. **C,** Contour-plots to monitor *in vitro* generation of the IRF4^hi^PAX5^low^ population; percentages are indicated. Bar-graph, percentages of the IRF4^hi^PAX5^low^ population from live cells. **D,** As in C but for the BLIMP1^+^PAX5^low^ population. Each dot corresponds to one mouse; mean±SEM is shown. Statistical analysis: two-tailed paired (A) and unpaired (C, D) parametric Student’s t-test and edgeR (B). *, *p*<0.05; **, *p*<0.01; ***, *p*<0.001.

### DGKζ promotes anabolism and cell cycle progression downstream of CD40 signaling

To understand how DGKζ might influence CD40-mediated B cell proliferation, we labeled WT and DGKζ^−/−^ B cells with CellTrace violet (CTV) fluorescent dye and measured the number of cell divisions by flow cytometry after a 96-h culture period with anti-CD40/IL-4/IL-5. DGKζ^−/−^ B cells exhibited a significant reduction (50%) in the frequency of cells undergoing five or more divisions (C5 region) compared with WT cells, with a concomitant increase in the number of non-dividing cells (C1 region) (Figure 3A). We then analyzed cell cycle distribution by propidium iodide staining at 48 h post-stimulation, when the first round of cell division was observed by CTV dilution (Figure S3A). DGKζ^−/−^ B cells showed impaired entry into the G2/M phase relative to WT cells (Figure 3B). To gain deeper insights into the transcriptional changes associated with these phenotypic differences, we performed gene set enrichment analysis (GSEA) on RNA-seq data collected at 48-h post-CD40 stimulation. This analysis revealed a downregulation of several cell cycle-related biological processes (MH: hallmark gene sets, Mouse MSigDB Collections) in DGKζ^−/−^ B cells (Figure 3C). Consistent with this, the expression of genes associated with cell cycle progression (*Mki67, Cdc20, Cdk1, Pcna)* was also lower in DGKζ^−/−^ B cells. Notably, the CD40-mediated induction of *Bcl2* expression and other pro-survival genes (*Bcl2a1a, Bcl2l1*) was higher in DGKζ^−/−^ B cells than in WT cells (Figure 3D). These observations suggest that DGKζ^−/−^ B cells are primarily arrested in their growth phase rather than experiencing more cell death.

**Figure 3.**
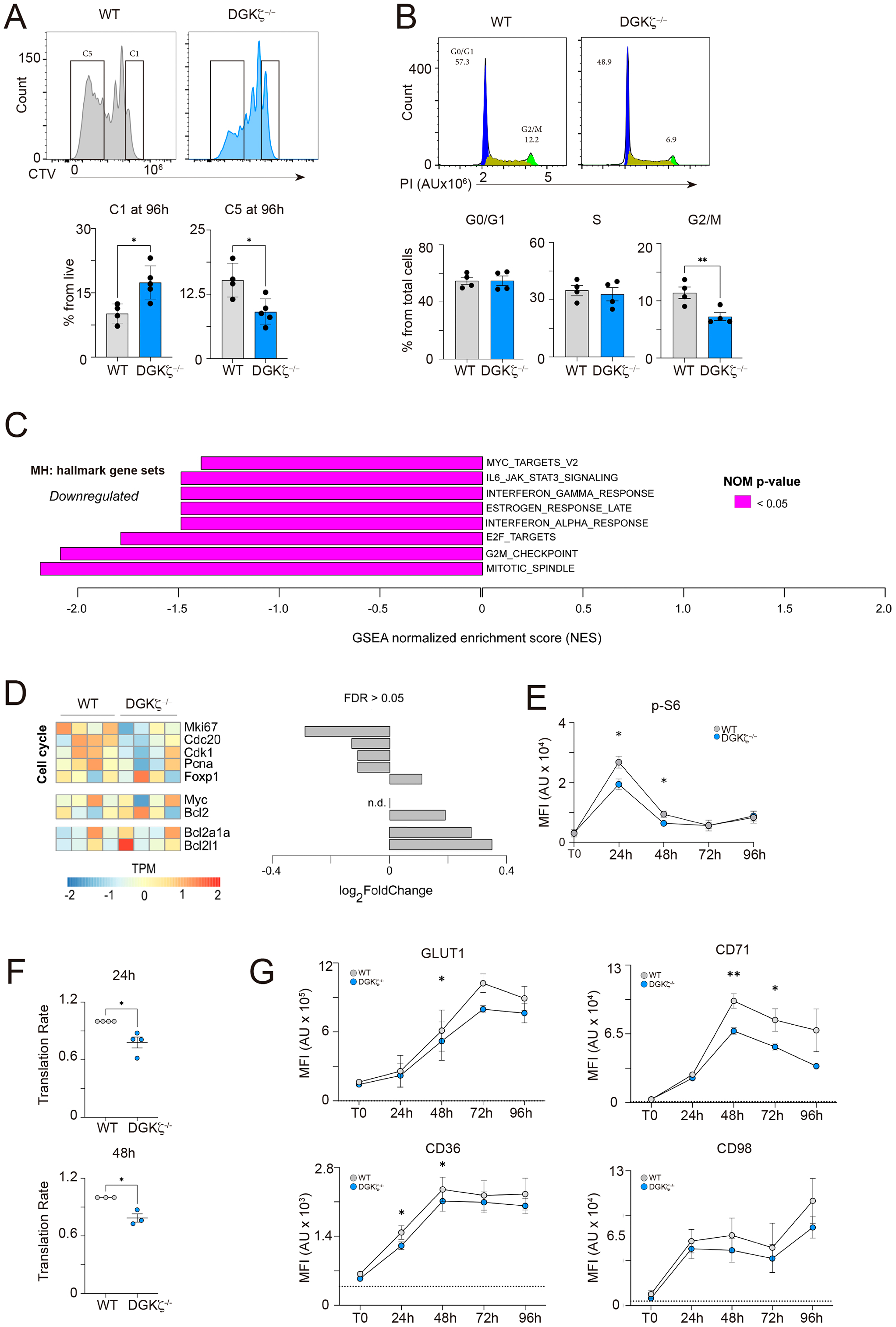
DGKζ promotes anabolism and cell proliferation downstream CD40 signaling. **A,** CTV-proliferation profiles of B cells cultured with anti-CD40/IL-4/IL-5 for 96 h, and percentages of the C1 (non-proliferated cells) and C5 (≥5 cell divisions) populations. **B,** B cells stimulated as in A for 48 h and collected for cell cycle analysis by propidium iodide (PI) staining. Cell cycle profiles (G0/G1 phase, blue; S phase, dark green; G2/M phase, light green) and percentages for each cell cycle phase. **C,** GSEA analysis using MH: hallmark gene sets, Mouse MSigDB Collections, for cell cycle related gene-sets in the RNA-seq data obtained from 48 h-stimulated B cells (n=4 mice per genotype). **D,** Heatmap and bar-plot illustrating the expression profiles (TPM, transcript per million mapped reads) and differential gene expression (log_2_FoldChange) of selected cell cycle-related genes obtained by RNA-seq, as in C (n=4 mice per genotype). **E-G**, B cells cultured as in A and analyzed by flow cytometry. **E**, Phosphorylated-S6 protein (p-S6) levels; MFI, mean fluorescence intensity; AU, arbitrary units (see Figure S3A). Data are the merge of n=5-6 mice per genotype. **F,** Protein translation rate measured by anti-puromycin staining of cells at 24- and 48-h post-stimulation. **G,** MFI values of the metabolite transporters GLUT1, CD71, CD98 and CD36 over time; data are the merge of n=3-5 mice per genotype. Each dot in A, B, F, corresponds to one mouse. Statistical analysis: two-tailed unpaired (A, E) and paired (B, F, G) parametric Student’s t-test, GSEA (C) and edgeR (D). *, *p*<0.05; **, *p*<0.01. Data in graphs are the mean ±SEM.

We monitored mTORC1 activity in cultures to assess the impact of DGKζ on cellular anabolism. We observed a peak of p-S6 at 24 h following CD40 stimulation in both WT and DGKζ^−/−^ B cells (Figures 3E and S3A). However, p-S6 levels were significantly lower in DGKζ^−/−^ B cells than in WT cells at 24 and 48 h (Figures 3E and S3A). To investigate protein synthesis, we conducted flow cytometry-based puromycin incorporation assays, finding that the protein translation rate was significantly lower in DGKζ^−/−^ B cells than in WT cells (Figure 3F), suggesting a compromised anabolic capacity. We also examined the expression of several metabolite transporters by flow cytometry, and consistently found lower expression of GLUT1, CD71, CD98 and CD36 in DGKζ^−/−^ B cells (Figures 3G and S3B). Therefore, the regulation of mTORC1 activation and metabolite transporter expression by DGKζ downstream of CD40 signaling may be crucial for the proliferation of B cells.

### DGKζ is essential for optimal mitochondrial function downstream of CD40 signaling in B cells

To investigate whether DGKζ deficiency affects mitochondrial remodeling, we monitored mitochondrial mass and membrane potential using MitoTracker Green FM and MitoTracker Red CMX ROS fluorescent probes, respectively (Figure S4A). Flow cytometry analysis revealed that mitochondrial mass was significantly lower in DGKζ^−/−^ B cells than in WT cells at baseline (Figure 4A). Following anti-CD40/IL-4/IL-5 stimulation, DGKζ^−/−^ B cells exhibited a trend towards decreased mitochondrial mass, although this difference did not reach statistical significance (Figure 4A). Notably, mitochondrial membrane potential was significantly lower in DGKζ^−/−^ B cells at 24 h, suggesting impaired mitochondrial function (Figure 4B). To further investigate this, we measured reactive oxygen species (ROS) production using CM-H2DCFDA and MitoSOX fluorescent probes, to estimate total and mitochondria-associated ROS production, respectively. We found that production of both total and mitochondrial-specific ROS was lower in DGKζ^−/−^ B cells than in WT cells at 48 h (Figures 4C-D and S4B), suggesting compromised mitochondrial function in the former.

**Figure 4.**
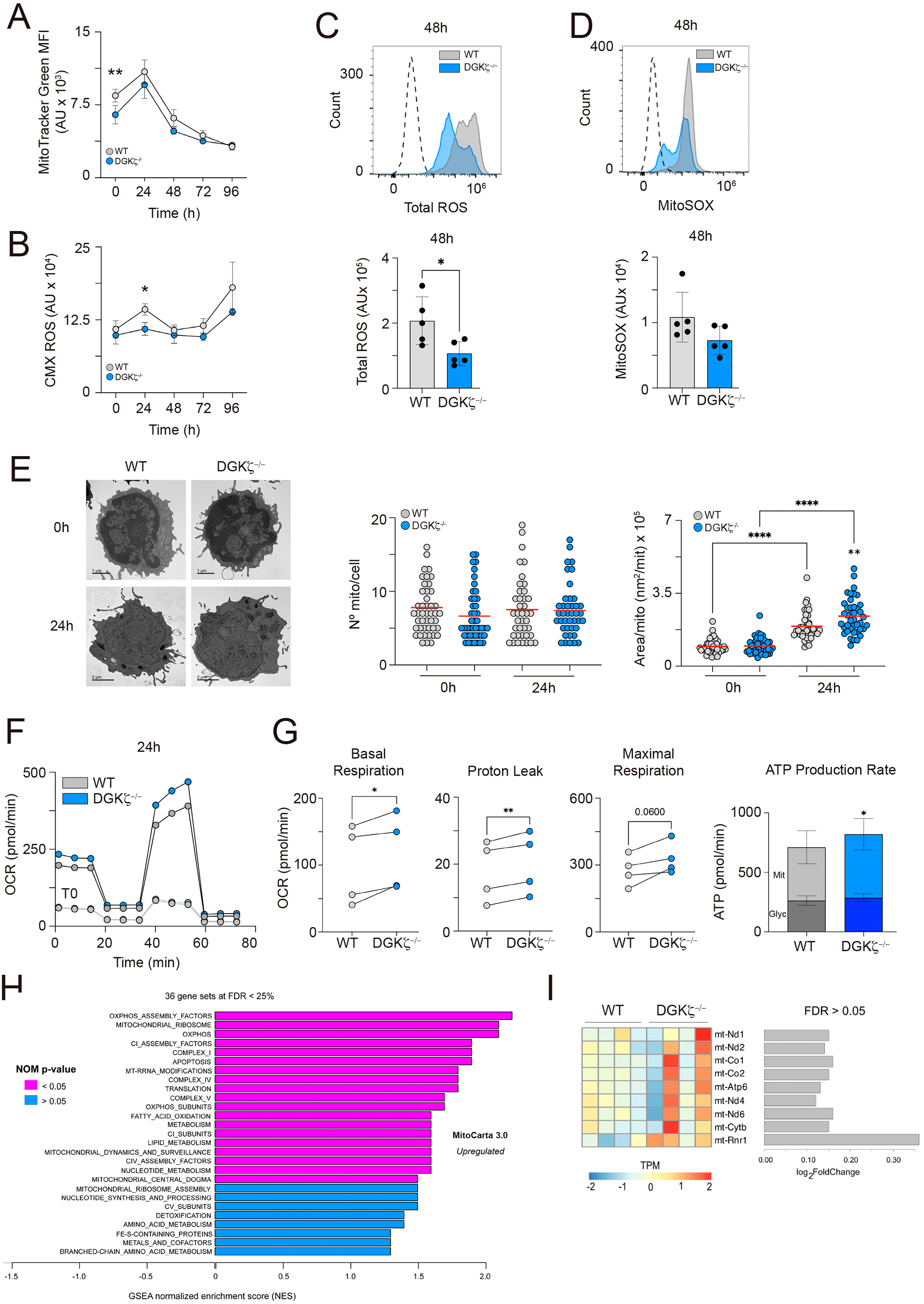
DGKζ is essential for optimal mitochondrial function downstream of CD40 signaling in B cells. **A-D,** B cells were stimulated with anti-CD40/IL-4/IL-5 and analyzed for mitochondrial parameters by flow cytometry. **A,** Mitochondrial mass (MitoTracker Green) and **B,** mitochondria membrane potential (CMXROS) measured over time. Mean Fluorescence Intensity (MFI) values in arbitrary units (AU) (see Figure S4A); data are the merge of n=5 mice per genotype. **C,** Profiles and MFI values per mouse of the total ROS content at 48 h. **D,** as in C but for mitochondrial ROS content (MitoSOX); dotted histogram, unstained cells. **E,** Mitochondrial content analysis of B cells freshly isolated (0h) and at 24-h post-stimulation. Electron microscopy images, quantification of the number of mitochondria (mito) per cell, and the mean-area per mitochondria in each cell are shown; each dot represents one cell (>40 cells analyzed per genotype and time point). Data are the merge of n=3 mice per genotype. **F-G,** Extracellular flux assays performed at the indicated time points (T0, freshly isolated; 24h post-stimulation). **F**, Profiles of oxygen consumption rate (OCR; in picomoles (pmol)/min) at T0 (grey line) and 24 h (black line). **G,** OCR values for parameters related to mitochondrial respiration (each dot corresponds to one mouse), and mean values of ATP production rate from glycolytic (Glyc; dark color) and mitochondrial (Mit; light color) pathways. **H,** GSEA analysis using MitoCarta3.0 of the RNA-seq data obtained from 48 h-stimulated WT and DGKζ^−/−^ B cells. **I,** Heatmap and bar-plot illustrating the expression profiles (TPM, transcript per million mapped reads) and differential gene expression (log_2_FoldChange) of selected mitochondrial genes obtained by RNA-seq, as in H. Data analysis in H, I, performed with n=4 mice per genotype. Statistical analysis: two-tailed paired parametric Student’s t-test (A-D, G), unpaired non-parametric Kolmogorov-Smirnov test (E, left graph), One-Way ANOVA (E, right graph), GSEA (H) and edgeR (I). Bar-graphs, mean±SEM. *, *p*<0.05; **, *p*<0.01.

Mitochondrial transcription factor A (TFAM) is a nuclear-encoded protein that regulates the copy number and transcription of mitochondrial DNA (mtDNA). The mRNA levels of *Tfam* were higher in DGKζ^−/−^ B cells than in WT cells following CD40 stimulation (Figure S4C). We detected increased TFAM protein levels in both genotypes following stimulation, as measured flow cytometry, with lower levels in DGKζ^−/−^ B cells (Figure S4D). To assess mitochondrial structure and content at the single-cell level, we performed electron microscopy at baseline and at 24 h post-stimulation (Figure 4E). Consistent with the MitoTracker Green analysis (Figure 4A), the number of mitochondria per cell was lower in unstimulated DGKζ^−/−^ B cells than in equivalent controls (Figure 4E). After 24 h, both genotypes showed an increase in mitochondrial number, with DGKζ^−/−^ cells reaching levels comparable with WT cells (Figure 4E). While not statistically significant, measurement of mtDNA copy number showed a similar trend (Figure S4G). Additionally, individual mitochondria were significantly larger in DGKζ^−/−^ B cells than in WT cells (Figure 4E). Overall, these results suggest that while mitochondria biogenesis may be slightly impaired in unstimulated DGKζ^−/−^ cells, it can be fully compensated for upon CD40 stimulation.

We next assessed mitochondrial OxPhos capacity at baseline and after CD40 stimulation using extracellular flux analysis. Comparison of the oxygen consumption rate at basal and maximal respiratory capacities revealed higher values for DGKζ^−/−^ cells than for WT cells at 24 h, which correlated with enhanced mitochondrial ATP production (Figures 4F-G). Contrastingly, glycolytic-associated ATP production remained comparable between both cell types (Figure 4G). Similar results were found at 48 h post-stimulation, albeit with less pronounced differences (Figures S4E-F). To investigate the underlying mechanism of the enhanced mitochondrial function in DGKζ^−/−^ B cells, we revisited the transcriptome data obtained at 48 h post-stimulation. GSEA for biological processes related to mitochondria metabolism (MitoCarta3.0^30^) revealed upregulation of numerous gene-sets in DGKζ^−/−^ B cells compared with controls (Figure 4H). These included genes involved in OxPhos and its assembly factors, mitochondrial ribosomes and mitochondrial dynamics. Additionally, we examined the expression of genes encoded by mtDNA, such as the subunits of NADH dehydrogenase (*mt-Nd1, mt-Nd2, mt-Nd6*), ATP synthase (*mt-Atp6*) and COX1 (*mt-Co1, mt-Co2*), and found higher levels in DGKζ^−/−^ B cells (Figure 4I), which was confirmed by immunoblotting (Figure S4H). Collectively, these findings suggest that DGKζ is essential for optimal mitochondrial function downstream of CD40 signaling in B cells. In the absence of DGKζ, mitochondria adapt their function by increasing OxPhos capacity and mitochondria biogenesis.

### DGKζ is needed for B cell progression in the germinal center response

We next performed T cell-dependent immunization experiments in mice to comprehensively analyze the germinal center response in DGKζ^−/−^ and WT B cells. We transferred CD45.2^+^ B cells (either WT or DGKζ^−/−^) from female spleens into CD45.1^+^ immunocompetent hosts (both female and male). One day later, these mice were immunized with NP-OVA antigen mixed with alum. Seven days post-immunization, we analyzed the immune response in the spleen and bone marrow using multi-parametric flow cytometry (Figure 5A).

**Figure 5.**
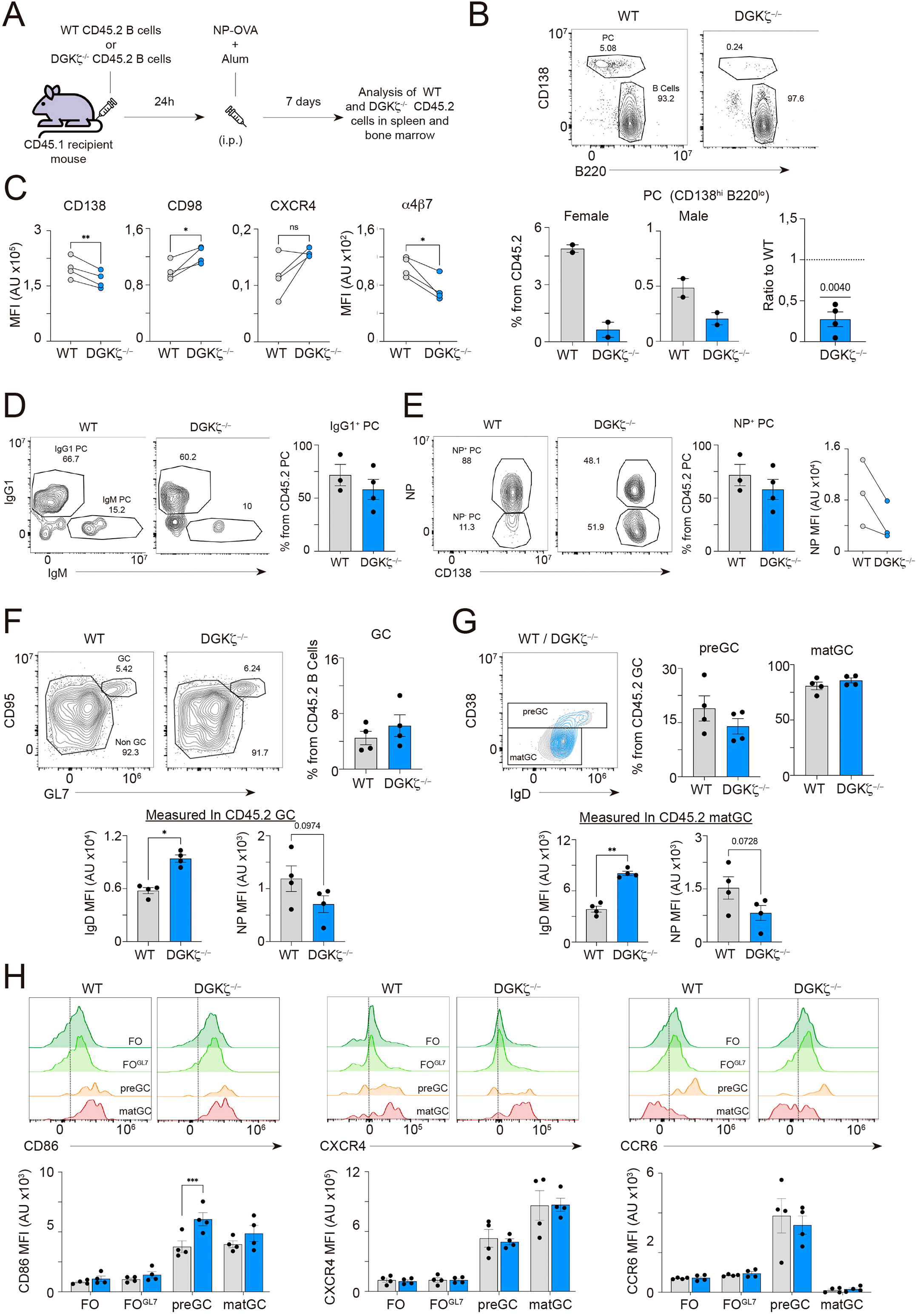
DGKζ is needed for B cell progression in the germinal center response. **A**, Experimental design for studying the *in vivo* response of CD45.2^+^ WT and DGKζ^−/−^ B cells in immunocompetent CD45.1^+^ host mice (see Figure S5A). **B**, Contour-plots of the CD45.2^+^ plasma cell population (PC; CD138^hi^B220^lo^; percentages are indicated). PC percentages for male and female, separately, and also normalized to CD45.2^+^ WT values. **C,** Mean fluorescence intensity (MFI) values of CD138, CD98, CXCR4, α4β7 for the CD45.2^+^ PC population. **D,** IgM/IgG1 contour-plots and IgG1^+^ PC percentages for the CD45.2^+^ PCs. **E,** NP-antigen/CD138 contour-plots in CD45.2^+^ PC. Graphs, frequencies of NP^+^ PC and MFI values of NP. **F,** GL7/CD95 contour-plots for gated CD45.2^+^ cells, indicating the germinal center (GC; GL7^+^CD95^+^) and non-GC (GL7^−^CD95^−^) populations; percentages are depicted. Right, GC percentages per mouse. Bottom, MFI values of IgD and NP in CD45.2^+^ GC B cells. **G,** Overlayed CD38/IgD counter-plots for WT (grey) and DGKζ^−/−^ (blue) GC B cells; gating of the pre-GC (CD38^+^IgD^+^) and mature-GC (CD38^−^ IgD^−^) populations is shown. Right, percentages of both populations. Bottom, MFI values of IgD and NP in CD45.2^+^ mature-GC B cells. **H,** Profiles and MFI values of CD86, CXCR4 and CCR6 in the follicular (FO), FO GL7^+^, pre-GC and mature-GC populations. Statistical analysis: one sample *t*-test (B) and two-tailed paired (C, F, G, H) and unpaired (D, E) parametric Student’s *t*-test. *, *p*<0.05; **, *p*<0.01; ***, *p*<0.001. Bar-graphs, mean± SEM; each dot corresponds to one mouse.

The frequency of CD45.2^+^ cells in total splenocytes was similar between WT- and DGKζ^−/−^-transferred mice (Figure S5A). We found higher numbers of the plasma cell population (CD45.2^+^CD138^+^B220^low/-^) in females than males, indicating a stronger immune response in females. Notably, DGKζ^−/−^ B cells generated significantly fewer plasma cells than WT CD45.2^+^ B cells (Figure 5B), consistent with previous findings from our group^28^. Further characterization revealed that DGKζ^−/−^ plasma cells expressed significantly lower levels of CD138 and α4β7 integrin than WT plasma cells, while CD98 expression was significantly higher (Figure 5C). The class-switched IgG1^+^ plasma cell population was diminished in DGKζ^−/−^ cells, although this did not reach statistical significance (Figure 5D). To assess antigen specificity, we stained plasma cells with fluorophore-conjugated NP antigen, finding that NP^+^ plasma cell numbers were lower in the DGKζ^−/−^ population than in the WT population together with lower staining levels, suggesting defects not only in plasma cell differentiation but also in affinity maturation (Figure 5E). Similar results were observed in the bone marrow plasma cell compartment (Figure S5B). No differences were found in the CD45.1^+^ plasma cell population generated in host mice transferred with WT B cells compared with those transferred with DGKζ^−/−^ B cells (Figures S5C-E). In contrast, the CD45.2^+^ MBCs (B220^+^CD95^−^GL7^−^IgD^−^CD38^+^) were increased in the DGKζ^−/−^ population (Figure S5F).

We found no differences in the generation of germinal center cells (CD45.2^+^B220^+^GL7^+^CD95^+^) between DGKζ^−/−^ and WT B cells. However, DGKζ^−/−^ germinal center cells displayed an immature phenotype, characterized by IgD^high^ NP^low^ staining (Figure 5F). To further investigate germinal center maturation, we analyzed CD38 and IgD expression, as described^18^. While the frequencies of pre-(CD38^+^IgD^+^) and mature-(CD38^−^IgD^low^) germinal center subpopulations were similar between the two genotypes, mature-germinal center DGKζ^−/−^ cells failed to downregulate IgD (Figure 5G). Notably, the host CD45.1^+^ germinal center population showed no differences in IgD levels or NP staining between DGKζ^−/−^- and WT-transferred mice (Figures S5G-H). We also examined the expression of other proteins related to germinal center maturation (CCR6, CXCR4, CD86) in these populations and at earlier stages: follicular (FO) B cells (CD45.2^+^B220^+^GL7^−^CD95^−^) and early activated FO B cells (CD45.2^+^B220^+^GL7^low^CD95^−^; FO^GL7^) (Figure S5F)^18^. CXCR4 and CCR6 upregulation from the FO^GL7^ to the pre-GC stage was similarly detected in both genotypes, with higher CD86 expression in DGKζ^−/−^ pre-germinal center cells than in WT cells (Figure 5H), indicative of increased activation. The transition from the pre- to the mature-germinal center stage is associated with CCR6 downregulation^18^, and this was likewise detected in DGKζ^−/−^ and WT cells (Figure 5H). No differences were observed in the centrocyte/centroblast populations (Figure S5J). Taken together, these findings indicate that DGKζ^−/−^ B cells can enter the germinal center pathway but experience impaired progression or blockage at the mature stage, reducing plasma cell generation.

### DGKζ drives F-actin polymerization and LFA-1-mediated adhesion at the CD40-mediated synapse-like contact

The CD40 signaling pathway is activated at the immune synapse, where antigen-experienced B cells initially interact with Tfh cells at the B/T border of the follicle. To investigate the molecular events triggered by CD40 stimulation within the B/T synapse and the role of DGKζ, we adapted our previous experimental model of artificial planar lipid bilayers used for BCR-mediated synapse studies^28^. We tethered anti-CD40 antibodies to the artificial membranes as surrogate CD40L (su-CD40L) *via* a biotin-streptavidin bridge (Figure S6A). To simulate physiological conditions, we estimated the density of CD40L molecules expressed at the surface of activated CD4^+^ T cells using calibrated beads, finding a range of CD40L expression up to 500 molecules/μm^2^ (Figure S6B). We subsequently employed two su-CD40L densities (100 and 500 molecules/μm^2^) on the artificial membranes. Using WT B cells, we monitored su-CD40L aggregation and the area of the established contact by confocal fluorescence microscopy and interference reflection microscopy (IRM), respectively (Figure S6C). Quantitative analysis revealed a direct correlation between su-CD40L density and the frequency of B cell contacts, their area and su-CD40L aggregation (Figure S6D). The incorporation of GPI-linked ICAM-1 adhesion molecules into the artificial membranes enhanced the contact area without affecting contact frequency or su-CD40L aggregation (Figure S6C, D). CD40 stimulation in B cells thus activates LFA-1 integrin, promoting adhesion through ICAM-1.

We next compared the behavior of DGKζ^−/−^ and WT B cells on the lipid bilayers containing ICAM-1 and tethered su-CD40L (Figure 6A). In the absence of DGKæ, B cells formed more and larger synaptic contacts (Figures 6B-C), with slightly increased su-CD40L aggregation (Figure 6D). Notably, half of the B cells analyzed from both genotypes exhibited plasma membrane ruffling during the recording period, suggesting enhanced actin remodeling in response to CD40 stimulation (Figure 6E; Movie S1). We fixed the B cells and examined F-actin filament distribution, observing a defined cluster of F-actin co-localizing with the su-CD40L aggregates, as well as F-actin-enriched peripheral structures (Figure 6F). Quantification of the F-actin detected at the contact plane revealed lower F-actin polymerization downstream of CD40 signaling in DGKζ^−/−^ B cells than in WT cells (Figure 6F). To further explore the specific contributions of DGKζ and its different domains in these events, we used A20 B cells transfected with the GFP-fused DGKζ constructs. Following CD40 stimulation, A20 cells displayed increased adhesion to the artificial membrane, accompanied by an increase in the size of synaptic contacts in the presence of ICAM-1, as found in primary B cells (Figure S6E). While ectopic expression of the DGKζ constructs had no effect on the frequency of cell-bilayer contacts (Figure S6F), it altered the shape and decreased the area of these contacts (Figures 6G-H). su-CD40L aggregation decreased with the expression of DGKζ-WT, but recovered with KD- or ΔPDZbm-mutant expression (Figure 6I). Time-lapse analysis of cell dynamics revealed a significant reduction in membrane ruffling and migratory behavior in DGKζ-WT-expressing cells compared with GFP^−^ control cells. This effect was less pronounced in cells expressing the KD- and ΔPDZbm-mutants (Figure 6J; Movie S2). These findings were corroborated by immunofluorescence analysis, which revealed decreased F-actin-enriched filopodia in DGKζ-WT-expressing cells, and a partial recovery in mutant-expressing cells (Figures S6G-H). Quantification of total F-actin accumulation located at the su-CD40L cluster showed no differences between DGKζ-WT or -KD expression when compared with GFP^−^ control cells, however, the ΔPDZbm-mutant significantly reduced F-actin accumulation (Figure S6I).

**Figure 6.**
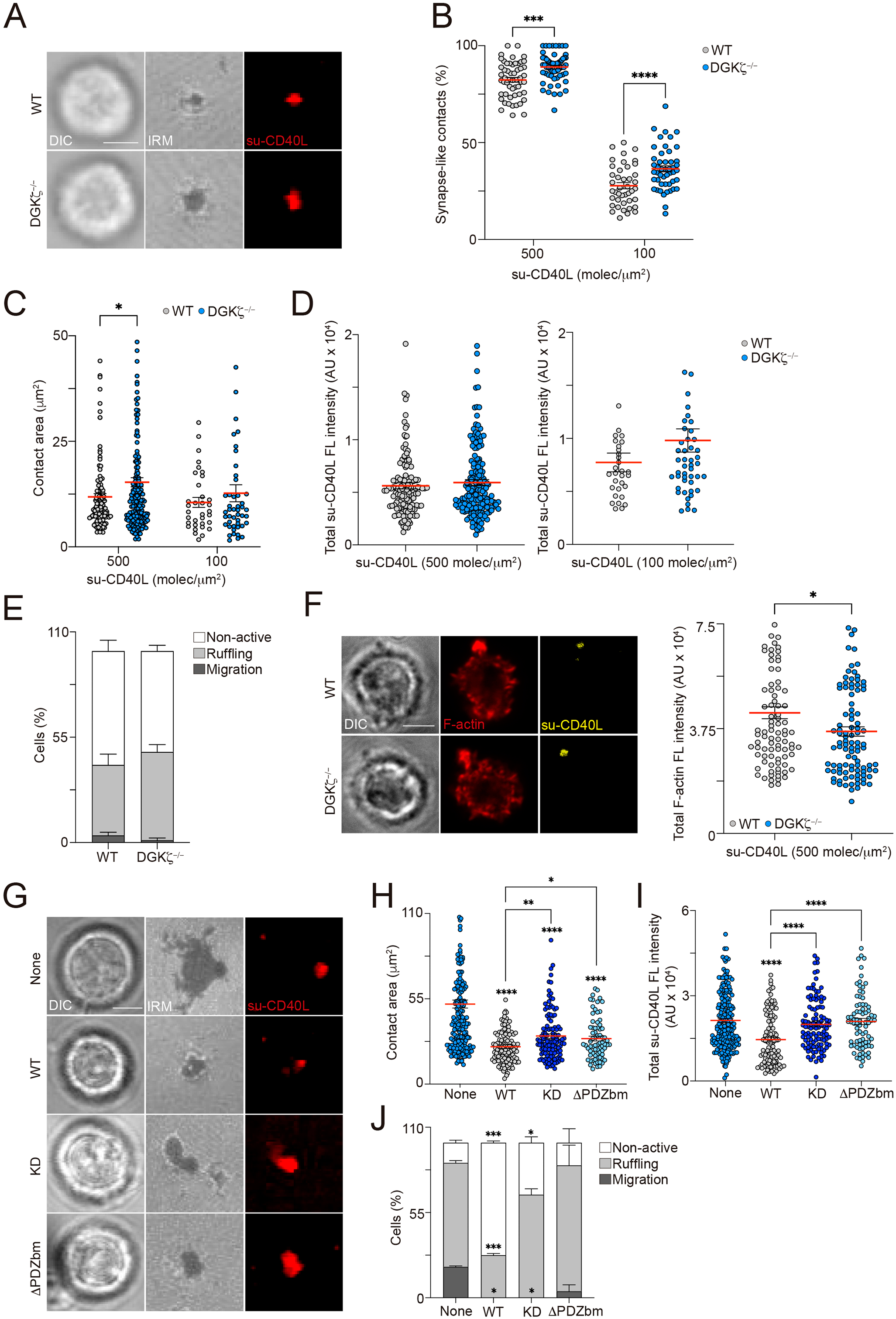
DGKζ drives F-actin polymerization and LFA-1-mediated adhesion at the CD40-mediated synapse-like contact. **A-F**, B cells were in contact with ICAM-1/su-CD40L-containing membranes (30 min, 37°C) and then imaged or fixed for immunofluorescence. **A**, DIC, IRM and fluorescence su-CD40L images at the contact plane of cells with a synapse-like contact (su-CD40L 500 molecules/μm^2^). **B**, Percentage of cells exhibiting a CD40-mediated synapse, estimated by IRM and fluorescence. **C**, Values of contact area (detected by IRM). **D**, Total su-CD40L fluorescence (FL) intensity, in arbitrary units (AU), at the synapse-like contact. **E**, Percentages of cells that showed migration (dark grey), were immobile but displaying membrane ruffling (light grey), or were immobile and without membrane ruffles (non-active; white). **F**, DIC, fluorescence F-actin and su-CD40L images of cells and quantification of the total F-actin FL intensity at the contact plane. **G-I**, A20 B cells transfected with the GFP-tagged WT, KD or ΔPDZbm DGKζ constructs assessed for CD40-mediated synapse formation, as above (su-CD40L 500 molecules/μm^2^). Electroporated cells negative for the expression of the construct (GFP^−^) were used as control (None). **G**, DIC, IRM, and fluorescence su-CD40L images at the contact plane of cells with a synapse-like contact. **H**, Values of contact area and **I**, of total su-CD40L FL intensity (in AU) at the synapse. **J**, as in E but for the specified A20 cells. Each dot in B corresponds to an imaged field, and to a single cell in the remaining. Data in B are the merged of n=6 experiments, in C, D, F corresponds to one representative experiment (n=6), and in E, H-J are the merge of n=3 experiments. Graphs, mean±SEM. Scale bar 2.5 μm. Statistical analysis: two-tailed unpaired (B, C, D) Student’s *t*-test, One-way (H, I) and two-way (E, J) ANOVA. *, *p*<0.05; **, *p*<0.01; ***, *p*<0.001; ****, *p*<0.0001.

Overall, the data indicate that CD40 engagement by su-CD40L tethered to the 2D, fluid surface triggers the formation of a CD40 cluster accompanied by an F-actin enriched domain beneath and surrounding LFA-1/ICAM-1 adhesions, resembling an immune synapse. Additionally, it induces the extension of exploratory membrane ruffles. DGKζ modulates these events in several ways: it elevates the threshold of CD40 stimulation needed to establish the synapse-like platform; it restricts the amount of CD40/su-CD40L accumulated at the cluster; and it diminishes the exploratory behavior of the B cell. These functions rely on both its kinase activity and its adaptor role through the PDZbm.

### CD40 stimulation drives microtubule-organizing center and organelle translocation to the immune synapse in a DGKζ-dependent manner

We next explored other molecular events, such as organelle translocation, that might accompany the establishment of the synapse driven by CD40 stimulation and contribute to its function. To simplify the visualization and quantification of organelle transport to the contact site, we employed 5-μm diameter streptavidin-coated silica beads, which mimic the size of T lymphocytes. These beads were either incubated with biotinylated-su-CD40L or left untreated as a control. We mixed these beads with B cells at a 1:1 ratio, allowed them to form conjugates for up to 30 min, fixed the cells, and performed immunofluorescence. Specifically, we stained the microtubule network and determined the position of the microtubule-organizing center (MTOC) in relation to the synapse-like contact by measuring distances (Figure S7A). The majority of WT cells exhibited MTOC translocation to the contact zone in response to CD40 stimulation, whereas DGKζ^−/−^ B cells did not (Figures 7A-B). We calculated a polarity index (PI) for each cell and categorized cells based on MTOC position (PI<0.4, near the contact; PI 0.4-0.8, in the mid-cell; PI>0.8, opposite the contact; Figure S7B). The distribution of DGKζ^−/−^ B cells based on PI values was similar for both uncoated and su-CD40L-coated beads, indicating that MTOC polarization in response to CD40 stimulation was impaired in these cells (Figure 7B). To further investigate this defect, we performed these assays in the presence of exogenous PA, finding that PA restored the ability of DGKζ^−/−^ B cells to translocate the MTOC to the synapse-like contact (Figure 7B). We also performed these experiments in A20 cells transfected with the different DGKζ constructs. Results showed that both the kinase activity and the PDZbm of DGKζ were essential for the optimal translocation of the MTOC to the CD40-mediated synapse contact (Figures S7C-D).

**Figure 7.**
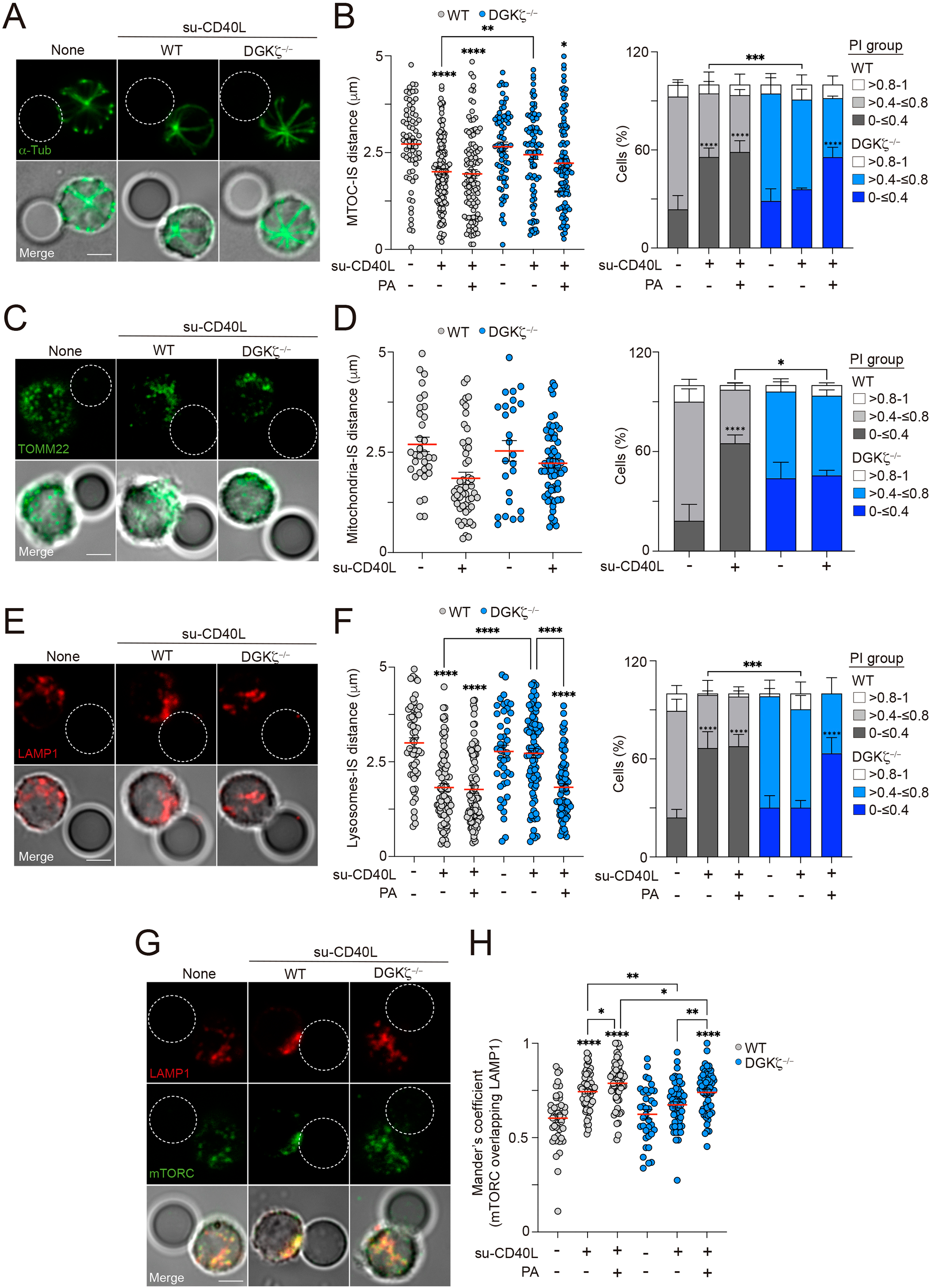
CD40 stimulation drives organelle translocation to the immune synapse in a DGKζ-dependent manner. B cells were mixed with unloaded (none) or su-CD40L-loaded beads in the absence or presence of phosphatidic acid (PA), and then fixed for immunofluorescence. **A**, Fluorescence alpha-tubulin (α-Tub) and DIC/fluorescence merge images of B cell-bead conjugates. White dashed-line circle, bead. **B**, Distance values of the MTOC to the synapse per cell and frequency of B cells distributed in each polarity index (PI) group (see Figure S7A, B). PI was calculated as the ratio of the MTOC-synapse distance and the diameter of the B cell. **C**, As in A but for TOMM22 staining, to detect mitochondria. **D**, As in B but for the mitochondria center-of-mass distance to the synapse and distribution in PI groups. **E**, As in A but for LAMP1 staining, to detect lysosomes. **F**, As in B but for the lysosomes center-of-mass distance to the synapse and distribution in PI groups. **G**, As in A but staining for lysosomes (LAMP1, red) and mTORC (green). **H**, Values of Mander’s coefficient for mTORC overlapping with LAMP1 (co-localization). Scale bar, 2.5 μm. Data shown in B, D, F and H corresponds to the merge of n=3 experiments; the mean±SEM is included. Statistical analysis: One-way (B, D, F, left graphs; H) and two-way (B, D, F, right graphs) ANOVA. *, *p*<0.05; **, *p*<0.01; ***, *p*<0.001; ****, *p*<0.0001.

Finally, we investigated the role of mitochondrial and lysosomal trafficking in the formation and function of su-CD40L-mediated synapse-like contacts. WT B cells demonstrated robust recruitment of mitochondria and lysosomes to the contact zone with su-CD40L-coated beads, as evidenced by immunofluorescence staining for TOMM22 (a component of the outer mitochondria membrane) and LAMP1 (lysosomal compartment), respectively, whereas DGKζ^−/−^ B cells exhibited impaired recruitment of both organelles (Figures 7C-F). Notably, exogenous PA supplementation restored lysosome trafficking in DGKζ^−/−^ B cells (Figure 7F). To explore the downstream consequences of impaired organelle trafficking, we focused on mTORC1 activation, which requires its recruitment from the cytosol to the lysosomal compartment. To question whether the reduced mTORC1 activation detected in DGKζ^−/−^ B cells downstream of CD40 signaling could be related to impaired mTORC1/lysosomal association, we co-stained the conjugates for mTORC1 and LAMP-1 and measured the co-localization coefficient. The mTORC1/lysosome co-localization values were higher for WT B cells forming conjugates with su-CD40L-coated beads than for those with uncoated-beads, and this was markedly reduced in DGKζ^−/−^ B cells (Figures 7G-H). Supplementation with exogenous PA restored mTORC1/lysosome co-localization in DGKζ^−/−^ B cells to levels resembling those of WT cells (Figure 7H). We also observed that PA increased the mTORC1/lysosome association rate in WT B cells stimulated with su-CD40L-coated beads (Figure 7H). These findings indicate that DGKζ is essential for the efficient trafficking of mitochondria, lysosomes and mTORC1 to the CD40-synapse. Its product, PA, appears to be important for these processes. The impaired lysosomal transport in DGKζ^−/−^ B cells likely contributes to the reduced mTORC1 activation observed upon CD40 stimulation. Overall, our results highlight the importance of DGKζ-mediated PA signaling in regulating cellular metabolism and immune responses through the coordinated trafficking of organelles and signaling molecules.

## DISCUSSION

Our data establish DGKζ as a crucial mediator of CD40 signaling in B cells by attenuating the activation of downstream effectors such as PKCβ and Akt. Concurrently, its local production of PA is essential for mTORC1 activation and organelle reorganization in response to CD40 signaling. PKCβ-deficient B cells exhibit impaired mTORC1 activation, mitochondrial remodeling and cell polarity, leading to muted germinal center responses and compromised plasma cell differentiation^13^. Given that PKCβ activates DGKζ through phosphorylation^25^, we propose that part of the PKCβ-deficient phenotype in B cells arises from the absence of DGKζ-pathway activation. The intermediate metabolite heme, which promotes plasma cell differentiation by inhibiting BACH2, is synthesized less efficiently under conditions of high ROS^13^. As DGKζ deficiency reduces CD40-induced ROS production in B cells, the heme/BACH2/Blimp-1 axis is likely unaffected and not responsible for the decreased plasma cell generation detected in DGKζ^−/−^ B cells. The PI3K/Akt axis is well established in mTORC1 activation downstream of CD40 signaling in B cells^20,31,32^. Despite increased Akt activation in CD40-stimulated DGKζ-deficient B cells, mTORC1 activation was dampened. These findings suggest that DGKζ is required to couple Akt/RHEB with mTORC1 activation, potentially through the regulation of CD40-synapse formation and polarized lysosome trafficking.

Our findings from PA-reconstitution assays and using the DGKζ-KD mutant indicate that DGKζ-derived PA is essential for optimal mTORC1 activation downstream of CD40 signaling in B cells. In addition to its kinase activity, its adaptor function through PDZbm is also required. DGKζ associates with sortin nexin 27 (SNX27) through a PDZbm/PDZ interaction^33^. SNX27 is an adaptor protein involved in intracellular trafficking and endosomal signaling through its association with the retromer complex. In T lymphocytes, SNX27 is crucial for polarized trafficking of endosomal compartments to the immune synapse, and its silencing leads to defects in cytoskeleton organization and DGKζ translocation to the synapse^34,35^. Impaired retromer function in B cells results in lysosomal dysfunction^36^. B cells deficient for SNX5, another SNX family member, exhibit compromised lysosomal integrity and function at the BCR-driven synapse, affecting antigen extraction^37^. We propose that, downstream of CD40 signaling, the DGKζ/SNX27 interaction positions DGKζ and PA production at endolysosomal membranes, facilitating mTORC1 activation.

CD40 stimulation in the absence of DGKζ significantly reduced plasma cell generation, primarily due to decreased expression of the genes *Irf4* and *Prdm1*. Reduced B cell proliferation, protein translation and mTORC1 activity in CD40-stimulated DGKζ-deficient B cells indicated impaired anabolism and cell growth. Additionally, mitochondria activation was weakened in DGKζ^−/−^ B cells. This contrasted with an increased in OxPhos capacity and mitochondrial-related gene expression, potentially representing a mechanism to address defects in anabolism and energy demands. B cell-specific TFAM-deficient mouse models have demonstrated that optimal mitochondrial function is crucial for B cells to participate in the germinal center response, and is also involved in actin cytoskeleton and lysosomal compartment remodeling^17,18^. *In vivo* analysis revealed that DGKζ-deficient B cells were stalled in the germinal center, exhibiting phenotypic characteristics (IgD^hi^ NP^low^) indicative of impaired progression and reduced plasma cell generation. We hypothesize that the compromised anabolism and mitochondrial activity of DGKζ-deficient B cells following CD40 stimulation, triggered by cognate interactions with Tfh cells, may underlie these defects.

Effective communication between B cells and Tfh cells is crucial for a successful germinal center response. This interaction involves the assembly of an immune synapse platform, where CD40 signaling plays an important role. Wataru Ise and colleagues demonstrated that the fate of plasma cells is influenced by the strength of CD40 signaling, and they proposed that more stable contacts with Tfh cells facilitate this^38^. Also, Zhixin Jing and colleagues demonstrated that CD40 signal strength is directly proportional to the peptide-MHC class II density expressed by B cells, and determines the magnitude of the germinal center expansion but not plasmablast differentiation^39^. B cell deficiency in DOCK8 impairs the stability of the B cell-Tfh conjugate, leading to inadequate costimulation and a subsequent failure of the humoral immune response^40^. Recent studies have shown that CD40 functions as a mechanoreceptor, receiving and sensing CD40L signals, with the magnitude of these forces influencing CD40 signaling and the B cell response^41^. The actin-myosin cytoskeleton was found to play an important role in these forces, and the authors discussed the implication of adhesion molecules. We observed that CD40 stimulation activates LFA-1 adhesion to ICAM-1, induces the formation of a su-CD40L cluster enriched in F-actin and promotes the generation of membrane ruffles. This su-CD40L cluster located usually at the edge of the B cell, combined with peripheral LFA-1-mediated adhesion and F-actin polymerization, suggests a motile or exploratory behavior in B cells upon CD40 stimulation. This aligns with *in vivo* observations of B-T conjugates at the follicular border, where B cells actively migrate carrying attached Tfh cells^5^. At the CD40-synapse, we found that DGKζ promotes actin polymerization primarily beneath the su-CD40L cluster while reducing F-actin-enriched membrane ruffles and peripheral LFA-1 adhesion that support cell exploratory behavior. These effects were dependent on both its kinase activity and its PDZbm adaptor functions. Additionally, DGKζ influences cell polarity events at the CD40-synapse: lysosome translocation facilitates the assembly of signaling platforms such as mTORC1, and mitochondria might provide the necessary energy. We propose that DGKζ acts as a sensor of CD40 signal quality, coordinating actin rearrangements, integrin clustering and cell polarity events for optimal CD40 mechanotransduction and signaling at the B-Tfh synapse. Insufficient CD40 signaling or defects in synapse assembly can hinder effective B cell integration of T cell help, ultimately impairing the germinal center response.

We found it intriguing that CD40-stimulated B cells deficient in DGKζ exhibited enhanced OxPhos capacity, an enlarged mitochondrial compartment and an upregulation of gene sets associated with mitochondrial metabolism. This could be a compensatory mechanism linked to the increased activation of PKCβ and Akt observed in those cells. Whether the accumulation of these molecular events in B cells paused at the progression of the germinal center has deleterious consequences and contributes to disease will need further investigation. The clinical targeting of DGKζ is currently being explored^42–44^. The primary rationale behind this approach is to enhance DAG signaling to overcome T cell immunosuppression in tumors. However, DGKζ deficiency has been shown to impair B cell immune response in mouse models, affecting both BCR^28^ and CD40 signaling (this study). Given that CD40 and DGKζ are also expressed in macrophages and dendritic cells, the impact of pharmacological DGKζ inhibition on CD40 functions in these cells requires further investigation. Therefore, when designing therapeutic strategies targeting DGKζ, it will be crucial to consider both its PA- and adaptor-related functions.

## RESOURCE AVAILABILITY

### Lead contact

Further information and request for resources and reagents should be directed to and will be fulfilled by the lead contact, Yolanda R. Carrasco.

### Materials availability

Materials needed to evaluate the conclusions in the paper are listed in the key resources table or in the Supplemental Data.

### Data and code availability

RNA-seq data have been deposited at Gene Expression Omnibus (GEO) under accession code GSE283915 and are publicly available as of the date of publication. Any additional information required to reanalyze the data reported in this paper is available from the lead contact upon request.

## ACKNOWLEDGEMENTS

We thank the Electron Microscopy Service (CNB, Universidad Autónoma, Madrid) for preparing samples (Epon embedding), obtaining ultrathin sections and TEM visualization; the Advanced Light Microscopy Service, the Flow Cytometry Service and the Animal Facility at the CNB for giving us technical support, and Dr. Kenneth McCreath for editorial assistance.

## AUTHORS CONTRIBUTIONS

A.F.B. and A.F.R designed parts of the study, performed experiments, analyzed the data and assisted in manuscript preparation. L.F.C. assisted in some experiment. T.P. performed the bioinformatic analysis of the RNAseq data. B.S.E., M.I.P., and N.M.M. assisted with metabolism experiments and data analysis, and provided input into the project. T.E, A.S.A. and S.C. assisted in mitochondrial data analysis, performed experiments of mtDNA and mitochondrial protein, and provided input into the project. R.J.S. performed experiments, analyzed the data and provided input into the project. Y.R.C. designed and supervised all aspects of the work and wrote the manuscript.

## SUPPLEMENTAL INFORMATION

Document S1. Figure S1-S7, Movies legends and Supplemental Methods.

Document S2. Excel file containing RNA-seq analysis data, too large to fit in a PDF, related to Figures 2B, 3C-D, 4H-I and S4C.

Movies S1 and S2

## STAR * Methods

### KEY RESOURCES TABLE

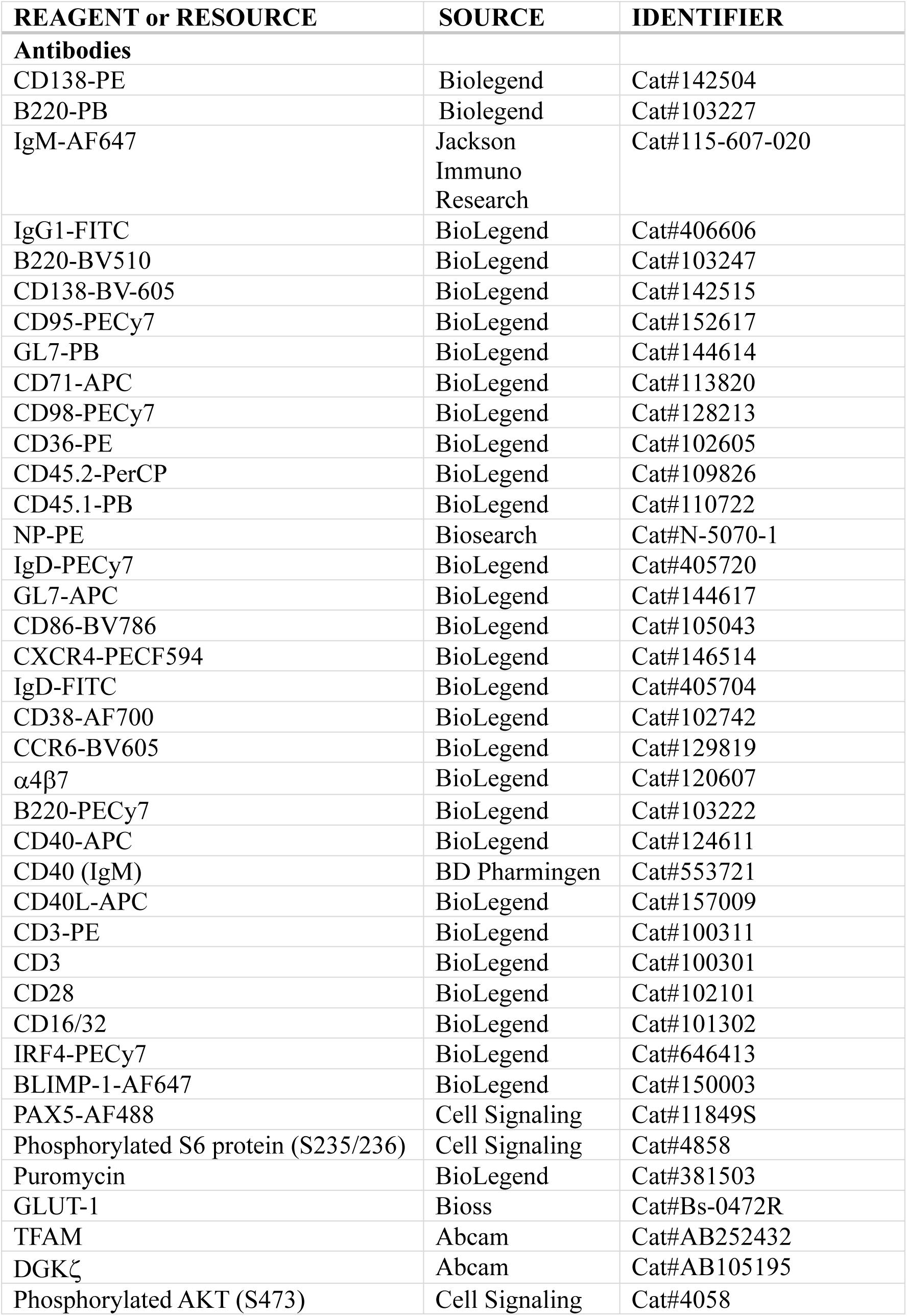
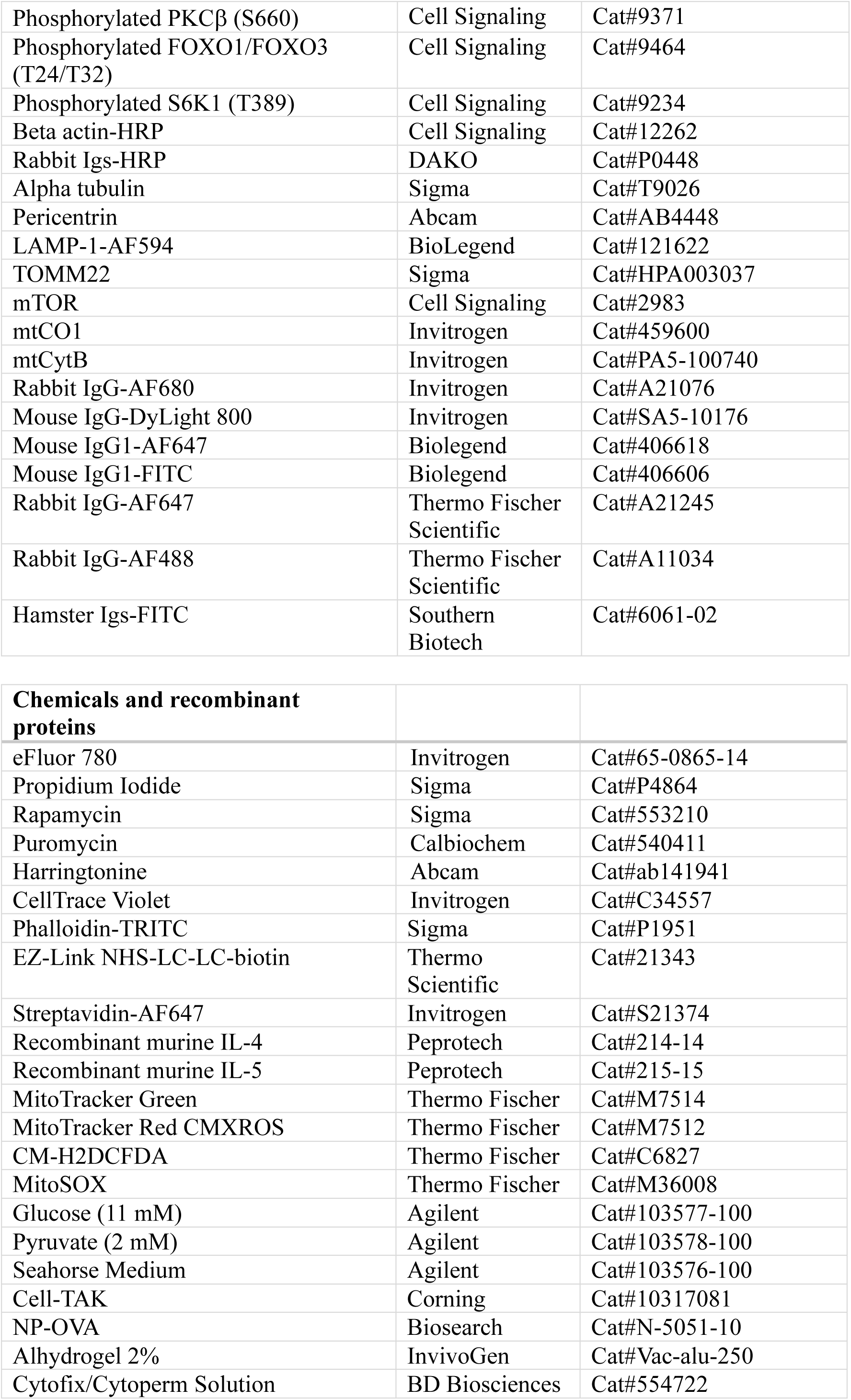
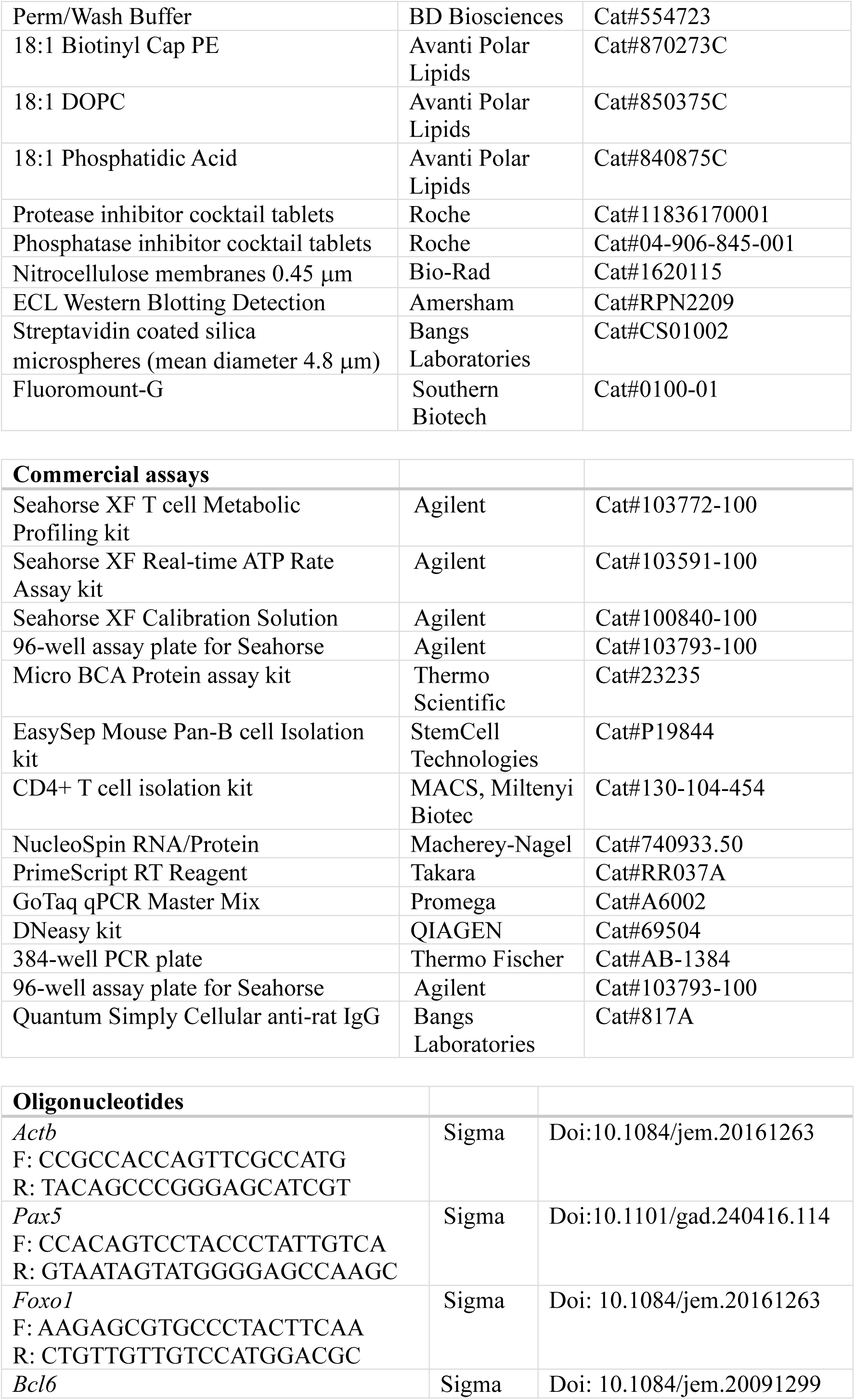
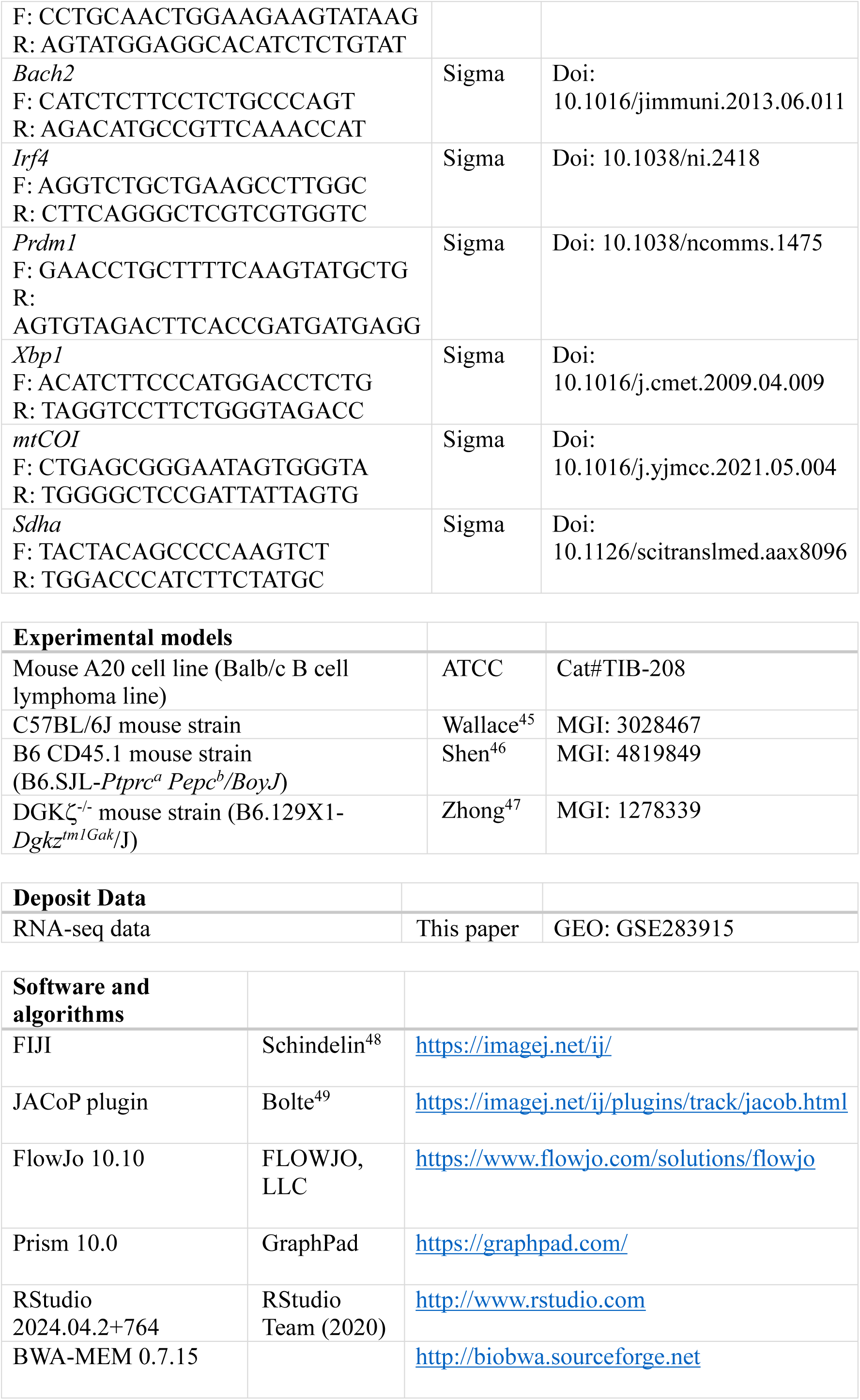
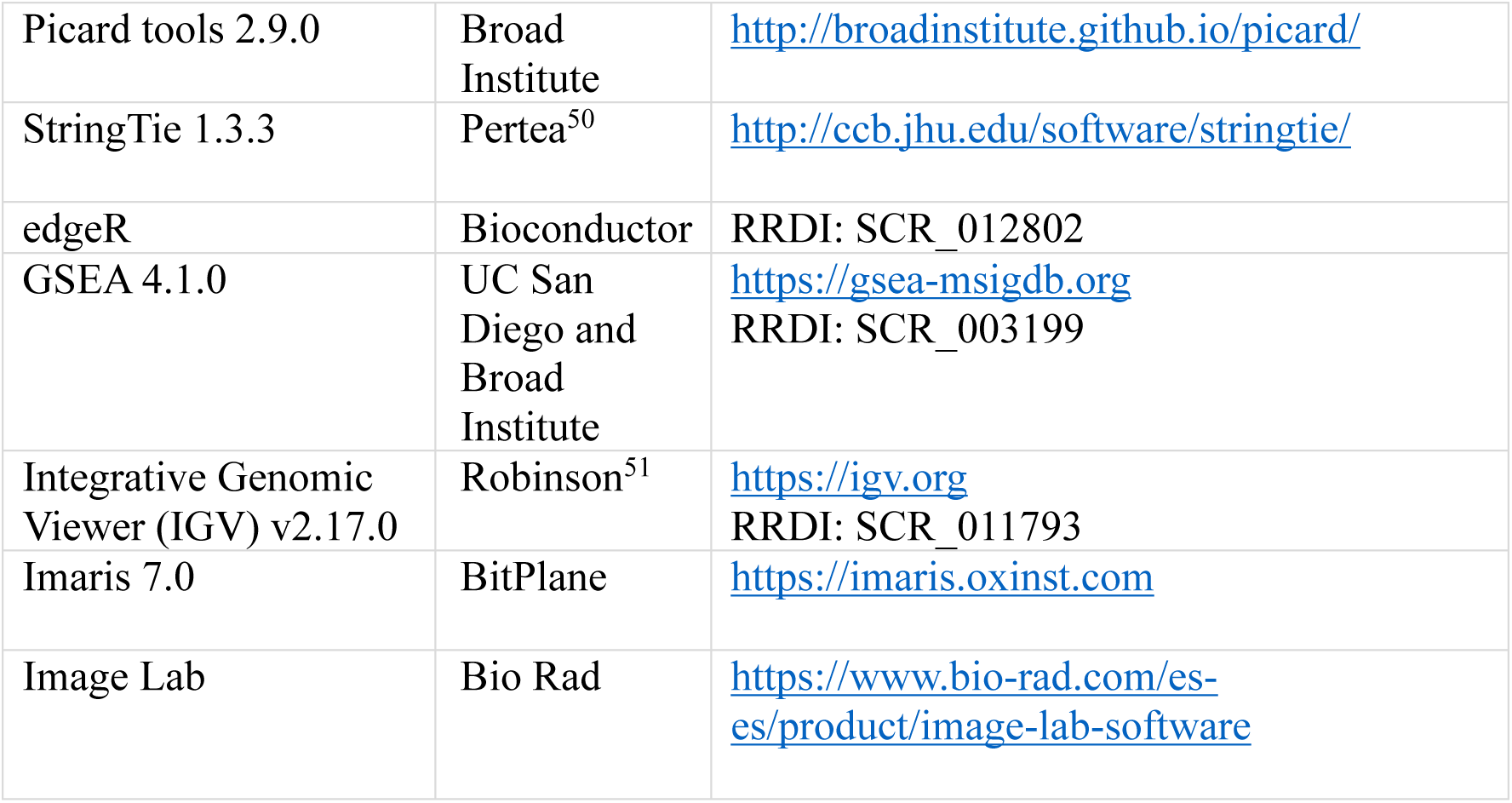

### METHODS

#### Mice and B cells

Primary B lymphocytes were isolated from the spleen of adult (4-7 months old) wild type (WT) and DGKζ^−/−^ mice^47^ on a C57BL/6 genetic background; males and females were used. Splenic B cells were purified by negative selection with EasySep^TM^ Mouse Pan-B cells Isolation Kit (StemCell Technologies); we enriched to > 95% B cells. CD4^+^ T cells were isolated from the spleen of adult WT mice by negative selection using a CD4^+^ T cell isolation kit (MACS, Miltenyi Biotec). Males and females of adult (3 months old) CD45.1^+^ mice on a C57BL/6 genetic background were used for in vivo experiments. Animal procedures were approved by the CNB-CSIC Bioethics Committee and conform to institutional, national and European Union (EU) regulations. Isolated B cells were resuspended in complete RPMI 1640 (cRPMI; 10 mM Hepes, 2 mM L-glutamine, 1 mM Na-Pyruvate, 50 μM β-mercaptoethanol, and 10,000 u/ml Penicillin/Streptomycin) supplemented with 10% complement-inactivated fetal calf serum (FCS) and either directly used or culture in p24 multi-well dishes (1.5×10^6^ cells/well and ml) in presence of hamster anti-mouse CD40 antibody (100 ng/ml), recombinant murine IL-4 and IL-5 (10 ng/ml each) for the indicated times. When required, B cells were labelled with CellTrace violet (CTV) to monitor cell proliferation by flow cytometry. Before culture, cells (5-10 × 10^6^ cells/ml) were incubated with 1 μM CTV in PBS (10 min, 37°C, in 15 ml falcon tubes), then a volume of FCS was added to stop the labelling reaction (1 min, room temperature, RT), the tube was filled with cRPMI 10% FCS for washing (440 g, 5 min, RT); cells were resuspended in the required medium volume. The A20 mouse B cell line was transiently transfected with plasmids encoding for GFP protein alone, or GFP-tagged DGKζ-WT, -kinase dead (KD) or -lacking the PDZ binding motif (ΔPDZbm) constructs^24^ by electroporation (260 mV, 950 μF) and were used 20 hours later. A20 transfected cells were cultured in cRPMI 10% FCS.

#### Western blotting

Freshly isolated primary B cells (15 × 10^6^) were cultured on a p35 plate in depletion medium (1.5 ml of cRPMI 0.5% FCS) for 1 h, split in 0.5 ml aliquots (5 × 10^6^ cells) and then stimulated with hamster anti-mouse CD40 antibody (100 ng/ml) for the indicated times. Cells were lysed in RIPA lysis buffer (50 mM Tris-HCl, pH 8.0, 150 mM NaCl, 1% NP40, 0.1% SDS, 0.5% Deoxycholate) with proteases and phosphatase inhibitors (Roche) (30 min, 4°C), lysates were centrifugated (20,000 g, 15 min, 4°C), and the supernatants were collected and stored at −80°C. Total protein concentration was quantified by Micro BCA Protein Assay Kit (Thermo Scientific). Proteins (10-20 μg/sample) were resolved by SDS-polyacrylamide gel electrophoresis (PAGE) and transferred to nitrocellulose membranes (Bio-Rad). Blots were blocked with TBS-T (10mM tris-HCl, pH 8.0, 150 mM NaCl, 0.1% Tween-20) 5% bovine serum albumin (BSA) (1 h, RT), incubated with the specified primary antibodies (overnight, 4°C), washed with TBS-T, incubated with horseradish peroxidase (HRP)-conjugated secondary antibody (1 h, RT), and finally washed with TBS-T. Signals were detected using the enhanced chemiluminescence (ECL) detection system (Amersham). Signal intensity values in arbitrary units (AU) for each protein were quantified with Image Lab software (Bio-Rad) and normalized to that of loading control (actin). For the detection of the mitochondrial proteins, the ODYSSEY Infrared Imaging System (LI-COR) was used.

#### Real-time microscopy on planar lipid bilayers

The artificial planar lipid bilayers were assembled in FCS2 chambers (Bioptechs) as described^52^. Briefly, unlabeled mouse GPI-linked ICAM-1 containing 1,2-dieloyl-PC (DOPC) liposomes and DOPC liposomes containing biotinylated lipids were mixed with DOPC liposomes at distinct ratios to obtain specific molecular densities (ICAM-1 at 200-300 molecules/μm^2^; biotin lipids as indicated in the figure legends). The artificial membranes (1 μl/membrane) were assembled on sulphocromic solution-treated coverslips, then blocked with phosphate-buffered saline (PBS)/2% FCS (1 h, RT), and incubated with Alexa Fluor 647-conjugated streptavidin (1:1000 dilution; 30 min, RT), followed by monobiotinylated hamster anti-mouse CD40 antibody (su-CD40L; 1:100 dilution; 30 min, RT). Monobiotinylation was achieved by labeling the antibody (0.5 mg/ml; 1 ml) with NHS-LC-LC-biotin (1 μg/ml, 30 min, RT, in PBS) followed by dialysis against PBS and analysis by flow cytometry. We estimated the number of molecules/μm^2^ of GPI-ICAM-1 or su-CD40L at the lipid bilayers by immunofluorometric assay with anti-ICAM-1 or anti-hamster-IgG antibodies, respectively. We obtained the standard values from microbeads with distinct calibrated IgG-binding capacities (Bangs Laboratories). Lipid stock in chloroform were obtained from Avanti Polar Lipids, Inc.

WT and DGKζ^−/−^ B cells (5 × 10^6^) were co-injected into the warmed chamber (37°C) for imaging. To distinguish them, one cell type was labeled with CTV (0.1 μM, 10 min, 37°C). Confocal fluorescence (1-μm optical section), DIC and IRM images were acquired. Consecutive videos were acquired when needed. Similarly, transfected A20 B cells (3 × 10^6^) were injected and imaged. Reagent dilution and cell assays were performed in chamber buffer (PBS, 0.5% FCS, 0.5 mg/ml D-glucose, 2 mM MgCl_2_ 0.5 mM CaCl_2_). Images were acquired on an Axiovert LSM 510 META inverted microscope with a 40x oil immersion objective (Zeiss) or STELLARIS 5 multispectral confocal system with a 63x oil immersion objective (Leica).

#### Immunofluorescence

Primary B cells or transfected A20 B cells (3-5 × 10^6^) were in contact with ICAM-1 lipid bilayers containing tethered su-CD40L for 30 min, fixed with 4% paraformaldehyde (PFA; 10 min, 37°C), permeabilized with PBS 0.1% Triton X-100 (5 min, RT), blocked with PBS 1% BSA, 0.1% goat serum, 0.05% Tween-20, 50 mM NaCl (overnight, 4°C), and stained with TRITC-conjugated phalloidin. For organelle translocation analysis, B cells and A20 B cells (2 × 10^6^) were mixed with un-loaded or su-CD40L-loaded streptavidin-coated beads (5 μm diameter; Bangs Laboratories) at a 1:1 ratio, incubated for 30 min at 37°C (Thermoblock, with shaking at 300 rpm) and then settled on poli-L-Lys coated slides (5 min) and fixed with 4% PFA (10 min, RT), permeabilized either with PBS 0.1% Triton X-100 for MTOC staining or PBS 0.05% Saponin 0.5% BSA for lysosomes and mitochondria staining (10 min, RT), blocked with PBS 1% BSA, 0.1% goat serum, 0.05% Tween-20, 50 mM NaCl or PBS 0.05% Saponin, 3% BSA, respectively (1 h, RT), and stained with anti-α-tubulin, anti-pericentrin, Alexa Fluor 594 conjugated anti-LAMP1, anti-TOMM22, or anti-mTOR followed by incubation with Alexa Fluor 647- or FITC-conjugated anti-mouse IgG1 or Alexa Fluor 647-conjugated anti-rabbit IgG. Slides were mounted using Fluoromount-G. For the exogenous PA assays, we used freshly prepared 10 mM PA stock (in 10 mM Tris-HCl, pH 8.0, 150 mM NaCl). The assays were performed in the presence of 0.1 mM PA during the stimulation time. FCS2 chamber and slides were imaged by confocal fluorescence microscopy as specified above.

#### Imaging data analysis

The frequency of synapse-like contact formation per imaged field was estimated as [number of B cells showing a su-CD40L cluster and IRM contact/total number of B cells (estimated by DIC)] × 100, using FiJi software. We used the IRM confocal image to focus on the B cell-artificial membrane contact and to define the synapse-like contact plane. Imaris 7.0 software was used for the quantitative analysis of fluorescence signals, as well as for cell contact area (IRM area). We set up the fluorescence background of the lipid bilayer for each su-CD40L density used. To measure cell behavior, we drew the area of the cell at time 0 (first frame) and then monitored membrane ruffling activity (plasma membrane extensions projected out of the drawn area) and migration (the cell moves out of the drawn area), using FiJi software. MTOC, lysosomes and mitochondria recruitment to the synapse-like contact in B cell-bead conjugates were quantified by projecting the distance between the center-of-mass of the fluorescent signal and the B cell-bead contact region (“a” distance) on the vector defined by the diameter of the cell that connect the contact region to the distal pole of the B cell (“b” distance) using the formula [“a” x cos-α] (see Fig S7A). The polarity index (PI) was calculated by dividing the projected distance [“a” x cos-α] between the diameter of the cell (“b”). The PI values range from 0 to 1, B cells were classified in three groups according to the PI value (PI<0.4, nearby the synapse contact; PI 0.4-0.8, in the mid-cell; PI>0.8, opposite to the synapse contact) (Fig S7B).

#### RT-qPCR and RNA-seq

RNA was isolated from naive and cultured B cells in presence of anti-CD40/IL-4/IL-5 for the indicated time points. The RNA extraction was performed using NucleoSpin RNA/Protein kit following the manufacturer instructions; purified RNA concentrations were measured using NanoDrop spectrophotometer. Retro-transcription was performed with PrimeScript RT Reagent Kit, using 500 ng of total RNA per sample. Quantitative PCR (qPCR) was performed to analyze each gene expression using the GoTaq qPCR Master Mix and QuantStudio™ 5 Real-Time PCR System (ThermoFisher); technical triplicates were done and the obtained values were normalized to actin expression as house-keeping gene. Primers used are described in the Key Resources Table. For transcriptome analysis using RNA-seq techniques, total RNA was isolated from B cells at 48 h after stimulation as mentioned. Libraries were prepared by the Genomics Unit of the Centro Nacional de Análisis Genómico (CNAG-CRG, Spain), and sequenced using NextSeq 2000 as paired-end 50-bp reads. Illumina FastQ files containing >30 million reads per sample were provided by the CNAG-CRG, who also carried out quality control of the FastQ files. For the data analysis, see Supplemental Methods.

#### Mitochondrial DNA content

Mitochondrial DNA content was measured as described^53^, with modifications. DNA extraction from B cell pellets was performed by DNeasy kit QIAGEN, and using 0.1 ml of elution buffer. Nuclear and mitochondrial DNA were amplified by quantitative PCR with 10 ng of total DNA as template, using specific primers for mitochondrial gene *mtCOI* and nuclear gene *SDHA*, and GoTaq qPCR Master mix. Mitochondrial content was normalized to nuclear DNA, and was calculated using the equation: ΔC_T_ = (nucDNA C_T_ – mtDNA C_T_). Relative mitochondrial DNA copies = 2 × 2^ΔCT^. The primers used are included in the Key Resources Table.

#### Adoptive cell transfer and Immunization

Freshly isolated CD45.2^+^ WT or DGKζ^−/−^ B cells (5 × 10^6^) were adoptively transferred to CD45.1^+^ immunocompetent mice by intravenous injection (>0.2 ml/mouse). After 24 h, the mice were immunized intraperitoneally with NP-OVA (100 ng) complexed with Alum (100 μl), in PBS (0.2 ml/mouse). Spleen and bone marrow were obtained seven days after immunization, processed, and total cell suspensions were treated with erythrocyte lysis buffer to eliminate erythrocytes. Finally, cells were resuspended in FACS buffer (PBS with 0.5% BSA, 2.5 mM EDTA) for staining and flow cytometry analysis. CD45.2 and CD45.1 populations were analyzed separately, and specific panels for plasma cells (CD138, B220, CXCR4, α4β7, CD98), class-switching (CD138, B220, IgM, IgD, IgG1) and germinal center populations (B220, GL7, CD95, IgD, CD38, CCR6, CD86 and CXCR4) were used.

#### Flow cytometry

We followed the protocols described in ^54^. In brief, cells were resuspended in FACS buffer and plated into p96-well conical-bottom plate. For membrane staining, 0.5 × 10^6^ cells (or 3 × 10^6^ for immunization experiments) were plated, incubated with unlabeled anti-CD16/32 antibody to prevent non-specific staining via Fc receptors (Fc-blocking; 10 min, 4°C) followed by addition of the antibody mix, including also efluor780 probe (1:2000 dilution) to exclude dead cells (30 min, 4°C). For intracellular staining 1 × 10^6^ cells were plated, incubated with Fc-blocking and efluor780 probe (15 min, 4°C), washed, and fixed with BD Cytofix/Cytoperm Fixation and Permeabilization Solution (20 min, 4°C). After washing with permeabilization buffer (BD Perm/Wash Buffer), cells were resuspended in the antibody mix prepared using the permeabilization buffer (30 min incubation, 4°C). When secondary antibodies were needed, cells were washed and resuspended in permeabilization buffer containing the secondary reagent (30 min incubation, 4°C). After staining, cells were washed, resuspended in FACS buffer and analyzed using a CytoFLEX flow cytometer (4 lasers; Beckman Coulter). FlowJo Software was used to analyze the acquired data.

For the analysis of mTORC1 activity at early time points, freshly isolated primary B cells (10 × 10^6^) were cultured on a p35 plate in depletion medium (1.5 ml of cRPMI 0.5% FCS; 1h), in presence of rapamycin (10 mM) when needed, then split in 0.5 ml aliquots (5 × 10^6^ cells) and stimulated with anti-CD40 antibody (100 ng/ml) and PA (0.1 mM) for the indicated time points. Then, intracellular staining was performed as above, using rabbit anti-pS6 antibody followed by Alexa Fluor 647-conjugated anti-rabbit secondary antibody. For intracellular TFAM detection, we followed the protocol described in ^17^. Briefly, cells were incubated with Fc-blocking and ef780 probe as above, then fixed with 4% PFA (15 min, RT), washed, and permeabilized with 90% freezer-cold methanol (10 min, RT). After washing with FACS buffer, anti-TFAM antibody was incubated in 50 μl FACS buffer supplemented with 2% goat serum (45 min, RT), followed by goat anti-rabbit secondary antibody (30 min, RT). For protein translation assay, cells were treated with puromycin (10 μg/ml; 45 min, 37°C); we used Harringtonine, inhibitor of protein translation, as a negative control. Then, cells were collected for intracellular staining using PE-conjugated anti-puromycin antibody (1 h, 4°C). To get the translation rate, the MFI values obtained for knock-out cells were normalized by those obtained for WT controls.

For the analysis of the cell cycle phases, cells were permeabilized with saponin 0.5% in FACS buffer (25 min, 4°C), washed, and stained with propidium iodide (50 μg/ml; 30 min, 4°C). After incubation, cells were directly analyzed at the flow cytometer. Data analysis performed with FlowJo Cell Cycle probabilistic analysis. For mitochondrial status assays, naive and CD40-stimulated B cells were incubated with the mitochondrial probes in RPMI 1640 (20 nM MitoTracker Green; 20 nM MitoTracker Red CMXROS; 1μM CM-H2DCFDA) or in HBSS medium (5μM MitoSox) for 30 min at 37°C, washed with FACS buffer, and analyzed.

For the quantification of CD40L expression in CD4^+^ T cells, a standard curve was generated using standard beads with known binding capacity for rat IgG molecules (Quantum^TM^ Simply Cellular anti-rat IgG; ABC; B=0, 1=4089, 2=20327, 3=98100, 4=243808). Beads were incubated with APC-conjugated rat anti-mouse CD40L antibody. Purified CD4^+^ T cells (4 × 10^5^) were cultured in flat-bottom p96 multi-well coated with anti-mouse CD3χ antibody (5 μg/ml) and in the presence of soluble anti-mouse CD28 antibody (0.5 μg/ml) for 20 h. Stimulated CD4^+^ T cells were stained with APC-conjugated rat anti-mouse CD40L. Flow cytometry of beads and cells was performed in a FACS Calibur cytometer. CD40L density was calculated using the equation 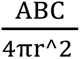.

#### Extracellular flux assays

Freshly isolated and CD40-activated B cells were resuspended in Seahorse medium, pH 7.4, supplemented with 11 mM Glucose and 2 mM Pyruvate (Agilent). Cells were seeded on a 96-well assay plate (Seahorse) coated with Cell-TAK (Corning). The assay was carried out on the XF96 Extracellular Flux analyzer. Oxygen Consumption Rates (OCR) were measured before and after the addition of different drugs included in the Seahorse XF Cell MitoStress kit and Seahorse XF Real-Time ATP Rate Assay kit.

#### Ultrastructural studies of mitochondria by electron microscopy

The assay was performed in collaboration with the Electron Microscopy Facility at the CNB-CSIC (Madrid, Spain). Naive and activated B cells were fixed in 2.5% glutaraldehyde 1% tannic acid (TAAB laboratories) in HEPES buffer 0.4 M pH 7.2 (Sigma-Aldrich; 2 h, RT), then washed with HEPES 0.4 M pH 7.2 (3 times, 15 min), post-fixed with 1 % osmium tetroxide (TAAB Laboratories) in potassium ferricyanide 0,8% (Sigma-Aldrich; 1 h, 4°C), and incubated with 2% aqueous uranyl acetate (Electron Microscopy Sciences; 1 h, 4°C). After washing with distilled water, samples were dehydrated with increasing concentrations of acetone (anhydrous, VWR) and embedded in epoxy resin TAAB 812 (TAAB Laboratories). Polymerization was carried out in epoxy resin 100% for 2 days, 60°C. Resin blocks were trimmed and ultrathin 70 nm-thick sections were obtained with the UC6 ultramicrotome (Leica Microsystems), transferred to 200 mesh nickel grids (Gilder) and stained with saturated aqueous uranyl acetate (20 min, RT) and lead citrate 0.2% (Electron Microscopy Sciences; 1 min, RT). Sections were visualized on a JEOL 1400 Flash electron microscope (operating at 100 kV). Micrographs were taken with a Gatan OneView digital camera at various magnifications. Image analysis was performed with FiJi software (NIH).

#### Statistical analysis

Graphs and statistical analyses were performed using Prism software (GraphPad). Two-tailed unpaired and paired Student’s t test, One-way ANOVA and Two-way ANOVA were applied. *, *p*<0.05; **, *p*<0.01; ***, *p*<0.001; ****, *p*<0.0001. For co-localization analysis, Mander’s overlap coefficients were calculated using the JACoP plugin (Just Another Co-localization plugin)^49^ of FIJI software. For RNA-seq data, edgeR was used. For the n° mitochondrias/cell, unpaired non-parametric Kolmogorov-Smirnov test was used.

## REFERENCES

1. Carrasco, Y.R., and Batista, F.D. (2007). B cells acquire particulate antigen in a macrophage-rich area at the boundary between the follicle and the subcapsular sinus of the lymph node. Immunity 27, 160–171. 10.1016/j.immuni.2007.06.007.

2. Phan, T.G., Grigorova, I., Okada, T., and Cyster, J.G. (2007). Subcapsular encounter and complement-dependent transport of immune complexes by lymph node B cells. Nat Immunol 8, 992–1000. 10.1038/ni1494.

3. Yuseff, M.I., Reversat, A., Lankar, D., Diaz, J., Fanget, I., Pierobon, P., Randrian, V., Larochette, N., Vascotto, F., Desdouets, C., et al. (2011). Polarized secretion of lysosomes at the B cell synapse couples antigen extraction to processing and presentation. Immunity 35, 361–374. 10.1016/j.immuni.2011.07.008.

4. Natkanski, E., Lee, W.Y., Mistry, B., Casal, A., Molloy, J.E., and Tolar, P. (2013). B cells use mechanical energy to discriminate antigen affinities. Science 340, 1587–1590. 10.1126/science.1237572.

5. Okada, T., Miller, M.J., Parker, I., Krummel, M.F., Neighbors, M., Hartley, S.B., O’Garra, A., Cahalan, M.D., and Cyster, J.G. (2005). Antigen-engaged B cells undergo chemotaxis toward the T zone and form motile conjugates with helper T cells. PLoS Biol 3, e150. 10.1371/journal.pbio.0030150.

6. Elgueta, R., Benson, M.J., de Vries, V.C., Wasiuk, A., Guo, Y., and Noelle, R.J. (2009). Molecular mechanism and function of CD40/CD40L engagement in the immune system. Immunol Rev 229, 152–172. 10.1111/j.1600-065X.2009.00782.x.

7. Ying, H., Li, Z., Yang, L., and Zhang, J. (2011). Syk mediates BCR- and CD40-signaling integration during B cell activation. Immunobiology 216, 566–570. 10.1016/j.imbio.2010.09.016.

8. Tang, T., Cheng, X., Truong, B., Sun, L., Yang, X., and Wang, H. (2021). Molecular basis and therapeutic implications of CD40/CD40L immune checkpoint. Pharmacol Ther 219, 107709. 10.1016/j.pharmthera.2020.107709.

9. Victora, G.D., and Nussenzweig, M.C. (2012). Germinal centers. Annu Rev Immunol 30, 429–457. 10.1146/annurev-immunol-020711-075032.

10. De Silva, N.S., and Klein, U. (2015). Dynamics of B cells in germinal centres. Nat Rev Immunol 15, 137–148. 10.1038/nri3804.

11. Roco, J.A., Mesin, L., Binder, S.C., Nefzger, C., Gonzalez-Figueroa, P., Canete, P.F., Ellyard, J., Shen, Q., Robert, P.A., Cappello, J., et al. (2019). Class-Switch Recombination Occurs Infrequently in Germinal Centers. Immunity 51, 337–350 e337. 10.1016/j.immuni.2019.07.001.

12. Ersching, J., Efeyan, A., Mesin, L., Jacobsen, J.T., Pasqual, G., Grabiner, B.C., Dominguez-Sola, D., Sabatini, D.M., and Victora, G.D. (2017). Germinal Center Selection and Affinity Maturation Require Dynamic Regulation of mTORC1 Kinase. Immunity 46, 1045–1058 e1046. 10.1016/j.immuni.2017.06.005.

13. Tsui, C., Martinez-Martin, N., Gaya, M., Maldonado, P., Llorian, M., Legrave, N.M., Rossi, M., MacRae, J.I., Cameron, A.J., Parker, P.J., et al. (2018). Protein Kinase C-beta Dictates B Cell Fate by Regulating Mitochondrial Remodeling, Metabolic Reprogramming, and Heme Biosynthesis. Immunity 48, 1144–1159 e1145. 10.1016/j.immuni.2018.04.031.

14. Gaudette, B.T., Jones, D.D., Bortnick, A., Argon, Y., and Allman, D. (2020). mTORC1 coordinates an immediate unfolded protein response-related transcriptome in activated B cells preceding antibody secretion. Nat Commun 11, 723. 10.1038/s41467-019-14032-1.

15. Jellusova, J. (2020). Metabolic control of B cell immune responses. Curr Opin Immunol 63, 21–28. 10.1016/j.coi.2019.11.002.

16. Urbanczyk, S., Baris, O.R., Hofmann, J., Taudte, R.V., Guegen, N., Golombek, F., Castiglione, K., Meng, X., Bozec, A., Thomas, J., et al. (2022). Mitochondrial respiration in B lymphocytes is essential for humoral immunity by controlling the flux of the TCA cycle. Cell Rep 39, 110912. 10.1016/j.celrep.2022.110912.

17. Yazicioglu, Y.F., Marin, E., Sandhu, C., Galiani, S., Raza, I.G.A., Ali, M., Kronsteiner, B., Compeer, E.B., Attar, M., Dunachie, S.J., et al. (2023). Dynamic mitochondrial transcription and translation in B cells control germinal center entry and lymphomagenesis. Nat Immunol 24, 991–1006. 10.1038/s41590-023-01484-3.

18. Iborra-Pernichi, M., Ruiz Garcia, J., Velasco de la Esperanza, M., Estrada, B.S., Bovolenta, E.R., Cifuentes, C., Prieto Carro, C., Gonzalez Martinez, T., Garcia-Consuegra, J., Rey-Stolle, M.F., et al. (2024). Defective mitochondria remodelling in B cells leads to an aged immune response. Nat Commun 15, 2569. 10.1038/s41467-024-46763-1.

19. Saxton, R.A., and Sabatini, D.M. (2017). mTOR Signaling in Growth, Metabolism, and Disease. Cell 168, 960–976. 10.1016/j.cell.2017.02.004.

20. Battaglioni, S., Benjamin, D., Walchli, M., Maier, T., and Hall, M.N. (2022). mTOR substrate phosphorylation in growth control. Cell 185, 1814–1836. 10.1016/j.cell.2022.04.013.

21. Foster, D.A. (2013). Phosphatidic acid and lipid-sensing by mTOR. Trends Endocrinol Metab 24, 272–278. 10.1016/j.tem.2013.02.003.

22. Avila-Flores, A., Santos, T., Rincon, E., and Merida, I. (2005). Modulation of the mammalian target of rapamycin pathway by diacylglycerol kinase-produced phosphatidic acid. J Biol Chem 280, 10091–10099. 10.1074/jbc.M412296200.

23. Torres-Ayuso, P., Tello-Lafoz, M., Merida, I., and Avila-Flores, A. (2015). Diacylglycerol kinase-zeta regulates mTORC1 and lipogenic metabolism in cancer cells through SREBP-1. Oncogenesis 4, e164. 10.1038/oncsis.2015.22.

24. Gharbi, S.I., Rincon, E., Avila-Flores, A., Torres-Ayuso, P., Almena, M., Cobos, M.A., Albar, J.P., and Merida, I. (2011). Diacylglycerol kinase zeta controls diacylglycerol metabolism at the immunological synapse. Mol Biol Cell 22, 4406–4414. 10.1091/mbc.E11-03-0247.

25. Rincon, E., Gharbi, S.I., Santos-Mendoza, T., and Merida, I. (2012). Diacylglycerol kinase zeta: at the crossroads of lipid signaling and protein complex organization. Prog Lipid Res 51, 1–10. 10.1016/j.plipres.2011.10.001.

26. Merida, I., Andrada, E., Gharbi, S.I., and Avila-Flores, A. (2015). Redundant and specialized roles for diacylglycerol kinases alpha and zeta in the control of T cell functions. Sci Signal 8, re6. 10.1126/scisignal.aaa0974.

27. Baldanzi, G., Bettio, V., Malacarne, V., and Graziani, A. (2016). Diacylglycerol Kinases: Shaping Diacylglycerol and Phosphatidic Acid Gradients to Control Cell Polarity. Front Cell Dev Biol 4, 140. 10.3389/fcell.2016.00140.

28. Merino-Cortes, S.V., Gardeta, S.R., Roman-Garcia, S., Martinez-Riano, A., Pineau, J., Liebana, R., Merida, I., Dumenil, A.L., Pierobon, P., Husson, J., et al. (2020). Diacylglycerol kinase zeta promotes actin cytoskeleton remodeling and mechanical forces at the B cell immune synapse. Sci Signal 13. 10.1126/scisignal.aaw8214.

29. Cook, S.L., Franke, M.C., Sievert, E.P., and Sciammas, R. (2020). A Synchronous IRF4-Dependent Gene Regulatory Network in B and Helper T Cells Orchestrating the Antibody Response. Trends Immunol 41, 614–628. 10.1016/j.it.2020.05.001.

30. Rath, S., Sharma, R., Gupta, R., Ast, T., Chan, C., Durham, T.J., Goodman, R.P., Grabarek, Z., Haas, M.E., Hung, W.H.W., et al. (2021). MitoCarta3.0: an updated mitochondrial proteome now with sub-organelle localization and pathway annotations. Nucleic Acids Res 49, D1541–D1547. 10.1093/nar/gkaa1011.

31. Sander, S., Chu, V.T., Yasuda, T., Franklin, A., Graf, R., Calado, D.P., Li, S., Imami, K., Selbach, M., Di Virgilio, M., et al. (2015). PI3 Kinase and FOXO1 Transcription Factor Activity Differentially Control B Cells in the Germinal Center Light and Dark Zones. Immunity 43, 1075–1086. 10.1016/j.immuni.2015.10.021.

32. Vivas-Garcia, Y., and Efeyan, A. (2022). The metabolic plasticity of B cells. Front Mol Biosci 9, 991188. 10.3389/fmolb.2022.991188.

33. Rincon, E., Santos, T., Avila-Flores, A., Albar, J.P., Lalioti, V., Lei, C., Hong, W., and Merida, I. (2007). Proteomics identification of sorting nexin 27 as a diacylglycerol kinase zeta-associated protein: new diacylglycerol kinase roles in endocytic recycling. Mol Cell Proteomics 6, 1073–1087. 10.1074/mcp.M700047-MCP200.

34. Gonzalez-Mancha, N., and Merida, I. (2020). Interplay Between SNX27 and DAG Metabolism in the Control of Trafficking and Signaling at the IS. Int J Mol Sci 21. 10.3390/ijms21124254.

35. Gonzalez-Mancha, N., Rodriguez-Rodriguez, C., Alcover, A., and Merida, I. (2021). Sorting Nexin 27 Enables MTOC and Secretory Machinery Translocation to the Immune Synapse. Front Immunol 12, 814570. 10.3389/fimmu.2021.814570.

36. Otomo, T., Schweizer, M., Kollmann, K., Schumacher, V., Muschol, N., Tolosa, E., Mittrucker, H.W., and Braulke, T. (2015). Mannose 6 phosphorylation of lysosomal enzymes controls B cell functions. J Cell Biol 208, 171–180. 10.1083/jcb.201407077.

37. Cabrera-Reyes, F., Contreras-Palacios, T., Ulloa, R., Jara-Wilde, J., Caballero, M., Quiroga, C., Feijoo, C.G., Diaz-Munoz, J., and Yuseff, M.I. (2025). SNX5 promotes antigen presentation in B cells by dual regulation of actin and lysosomal dynamics. Life Sci Alliance 8. 10.26508/lsa.202402917.

38. Ise, W., Fujii, K., Shiroguchi, K., Ito, A., Kometani, K., Takeda, K., Kawakami, E., Yamashita, K., Suzuki, K., Okada, T., and Kurosaki, T. (2018). T Follicular Helper Cell-Germinal Center B Cell Interaction Strength Regulates Entry into Plasma Cell or Recycling Germinal Center Cell Fate. Immunity 48, 702–715 e704. 10.1016/j.immuni.2018.03.027.

39. Jing, Z., McCarron, M.J., Dustin, M.L., and Fooksman, D.R. (2022). Germinal center expansion but not plasmablast differentiation is proportional to peptide-MHCII density via CD40-CD40L signaling strength. Cell Rep 39, 110763. 10.1016/j.celrep.2022.110763.

40. Deobagkar-Lele, M., Crawford, G., Crockford, T.L., Back, J., Hodgson, R., Bhandari, A., Bull, K.R., and Cornall, R.J. (2024). B cells require DOCK8 to elicit and integrate T cell help when antigen is limiting. Sci Immunol 9, eadd4874. 10.1126/sciimmunol.add4874.

41. Choi, H.K., Travaglino, S., Munchhalfen, M., Gorg, R., Zhong, Z., Lyu, J., Reyes-Aguilar, D.M., Wienands, J., Singh, A., and Zhu, C. (2024). Mechanotransduction governs CD40 function and underlies X-linked hyper-IgM syndrome. Sci Adv 10, eadl5815. 10.1126/sciadv.adl5815.

42. Wichroski, M., Benci, J., Liu, S.Q., Chupak, L., Fang, J., Cao, C., Wang, C., Onorato, J., Qiu, H., Shan, Y., et al. (2023). DGKalpha/zeta inhibitors combine with PD-1 checkpoint therapy to promote T cell-mediated antitumor immunity. Sci Transl Med 15, eadh1892. 10.1126/scitranslmed.adh1892.

43. Kureshi, R., Bello, E., Kureshi, C.T.S., Walsh, M.J., Lippert, V., Hoffman, M.T., Dougan, M., Longmire, T., Wichroski, M., and Dougan, S.K. (2023). DGKalpha/zeta inhibition lowers the TCR affinity threshold and potentiates antitumor immunity. Sci Adv 9, eadk1853. 10.1126/sciadv.adk1853.

44. Martin-Salgado, M., Ochoa-Echeverria, A., and Merida, I. (2024). Diacylglycerol kinases: A look into the future of immunotherapy. Adv Biol Regul 91, 100999. 10.1016/j.jbior.2023.100999.

45. Wallace, M.E. (1950). Locus of the gene ‘fidget’ in the house mouse. Nature 166, 407. 10.1038/166407a0.

46. Shen, F.W., Saga, Y., Litman, G., Freeman, G., Tung, J.S., Cantor, H., and Boyse, E.A. (1985). Cloning of Ly-5 cDNA. Proc Natl Acad Sci U S A 82, 7360–7363. 10.1073/pnas.82.21.7360.

47. Zhong, X.P., Hainey, E.A., Olenchock, B.A., Jordan, M.S., Maltzman, J.S., Nichols, K.E., Shen, H., and Koretzky, G.A. (2003). Enhanced T cell responses due to diacylglycerol kinase zeta deficiency. Nat Immunol 4, 882–890. 10.1038/ni958.

48. Schindelin, J., Arganda-Carreras, I., Frise, E., Kaynig, V., Longair, M., Pietzsch, T., Preibisch, S., Rueden, C., Saalfeld, S., Schmid, B., et al. (2012). Fiji: an open-source platform for biological-image analysis. Nat Methods 9, 676–682. 10.1038/nmeth.2019.

49. Bolte, S., and Cordelieres, F.P. (2006). A guided tour into subcellular colocalization analysis in light microscopy. J Microsc 224, 213–232. 10.1111/j.1365-2818.2006.01706.x.

50. Pertea, M., Pertea, G.M., Antonescu, C.M., Chang, T.C., Mendell, J.T., and Salzberg, S.L. (2015). StringTie enables improved reconstruction of a transcriptome from RNA-seq reads. Nat Biotechnol 33, 290–295. 10.1038/nbt.3122.

51. Robinson, J.T., Thorvaldsdottir, H., Winckler, W., Guttman, M., Lander, E.S., Getz, G., and Mesirov, J.P. (2011). Integrative genomics viewer. Nat Biotechnol 29, 24–26. 10.1038/nbt.1754.

52. Saez de Guinoa, J., Barrio, L., Mellado, M., and Carrasco, Y.R. (2011). CXCL13/CXCR5 signaling enhances BCR-triggered B-cell activation by shaping cell dynamics. Blood 118, 1560–1569. 10.1182/blood-2011-01-332106.

53. Cao, Y., Vergnes, L., Wang, Y.C., Pan, C., Chella Krishnan, K., Moore, T.M., Rosa-Garrido, M., Kimball, T.H., Zhou, Z., Charugundla, S., et al. (2022). Sex differences in heart mitochondria regulate diastolic dysfunction. Nat Commun 13, 3850. 10.1038/s41467-022-31544-5.

54. Jimenez-Saiz, R., Chu, D.K., Mandur, T.S., Walker, T.D., Gordon, M.E., Chaudhary, R., Koenig, J., Saliba, S., Galipeau, H.J., Utley, A., et al. (2017). Lifelong memory responses perpetuate humoral T(H)2 immunity and anaphylaxis in food allergy. J Allergy Clin Immunol 140, 1604–1615 e1605. 10.1016/j.jaci.2017.01.018.

