## Supplemental Data for "Diacylglycerol kinase ζ dictates CD40-mediated immune synapse formation, mTORC1 signaling and plasma cell fate in B lymphocytes"

### SUPPLEMENTAL FIGURES

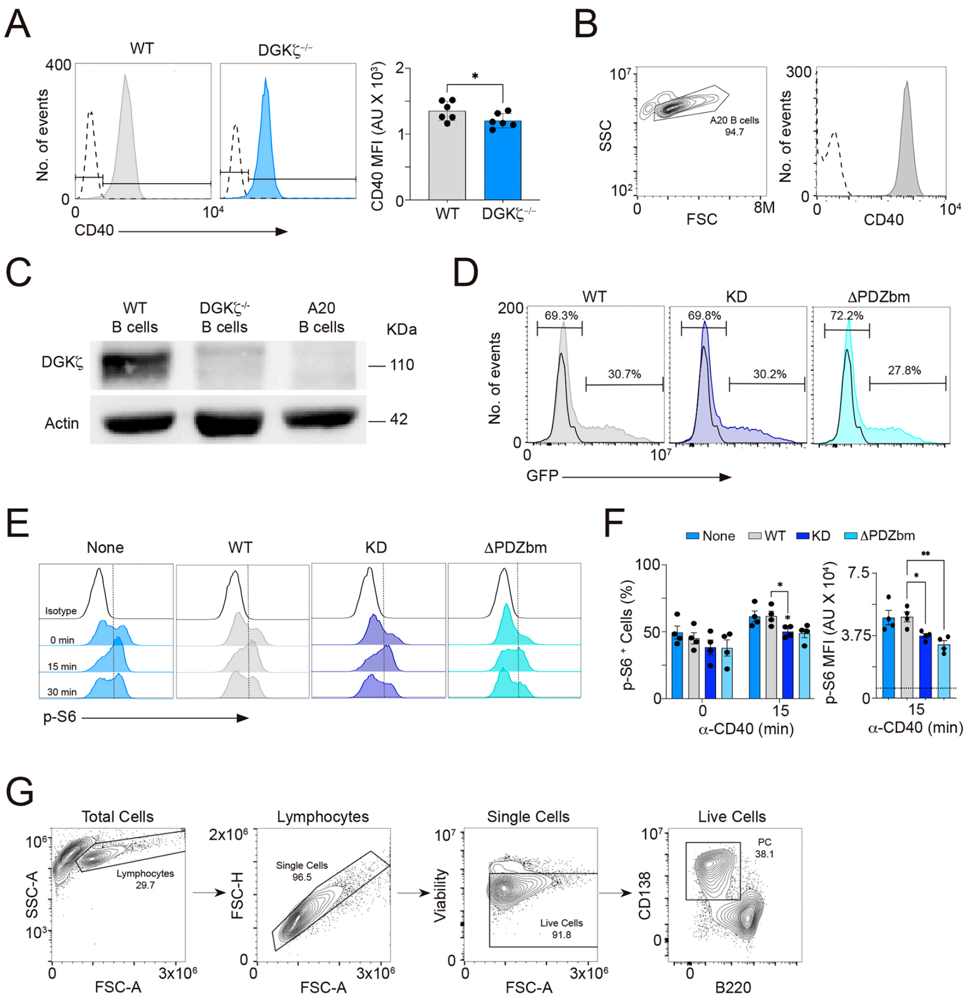


**Figure S1. DGKζ promotes mTORC1 activity and plasma cell generation downstream CD40. A,** Representative profiles of surface CD40 expression in naive WT and DGKζ$^{-/-}$ B cells; dotted line, isotype control. CD40 mean fluorescence intensity (MFI) values, in arbitrary units (AU), per mouse are shown. **B**, FSC/SSC contour-plot and surface CD40 expression profile in A20 B cells; dotted line, isotype control. **C**, Western Blot analysis of DGKζ expression in primary WT, DGKζ$^{-/-}$ B cells and A20 B cell line; actin, loading control. **D**, Profiles of GFP expression for A20 B cells transfected with DGKζ-WT, -kinase dead (KD) or -ΔPDZbm contructs (filled histograms), 20 h post-electroporation; percentages of GFP-negative and GFP-positive cells are indicated; black line histogram, non-transfected A20 cells. **E-F**, Transfected A20 B cells were stimulated with anti-mouse CD40 antibody for the specified time periods, then fixed, permeabilized and stained for the phosphorylated form of the S6 protein (p-S6). **E**, Profiles of p-S6 in gated GFP$^{-}$ cells (None; used as control) and GFP$^{+}$ cells for the three DGKζ constructs.

**F**, Quantification of the frequency of p-S6$^{+}$ cells (left) and values of p-S6 MFI (right; dotted line, isotype mean value) for the transfected A20 B cells. Data are the merge of n=4 experiments; mean ±SEM is shown. Statistical analysis: two-tailed unpaired Student's *t*-test and two-way ANOVA. *, $p$<0.05; **, $p$<0.01. **G**, Gating strategy used for the analysis of the plasma cells population (PC, CD138$^{hi}$ B220$^{low/-}$) generated at 96h.

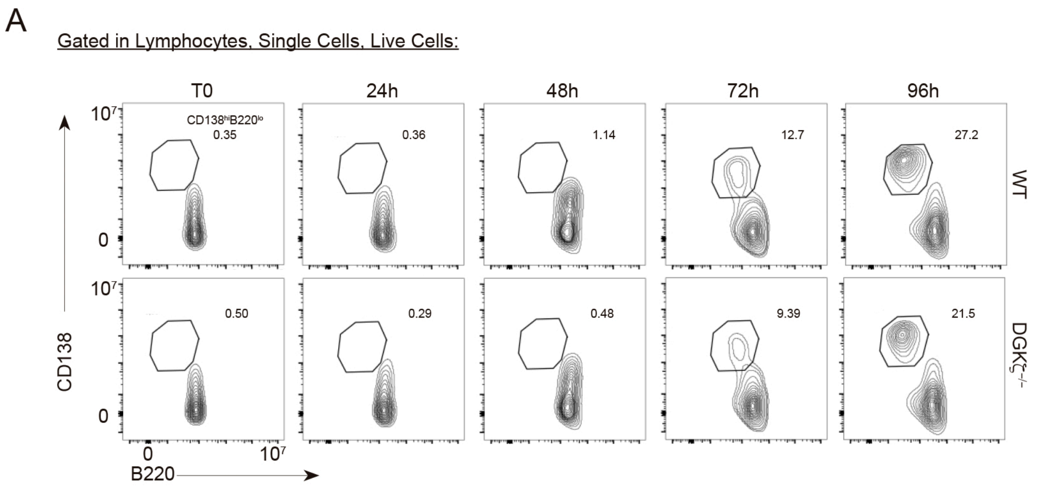


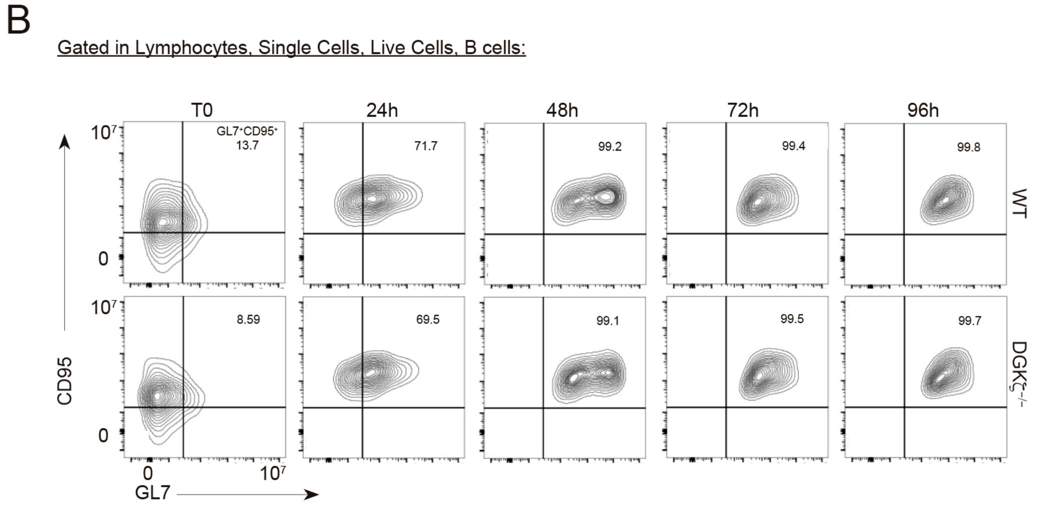


**Figure S2. Analysis of B220, CD138, CD95 and GL7 surface markers during the in vitro assays.** WT and DGK$\zeta^{-/-}$ B cells were cultured in the presence of anti-CD40/IL-4/IL-5 for different time points, collected and analyzed by flow cytometry. **A,** Representative CD138/B220 contour-plots; T0, freshly isolated B cells. The plasma cell (PC; CD138$^{hi}$ B220$^{lo}$) gate is shown, indicating the percentage at each time point. **B,** As in A but for GL7/CD95 markers; the generation of a germinal center-like (GC; GL7$^{+}$ CD95$^{+}$) population is followed, indicating the percentage at each time point.

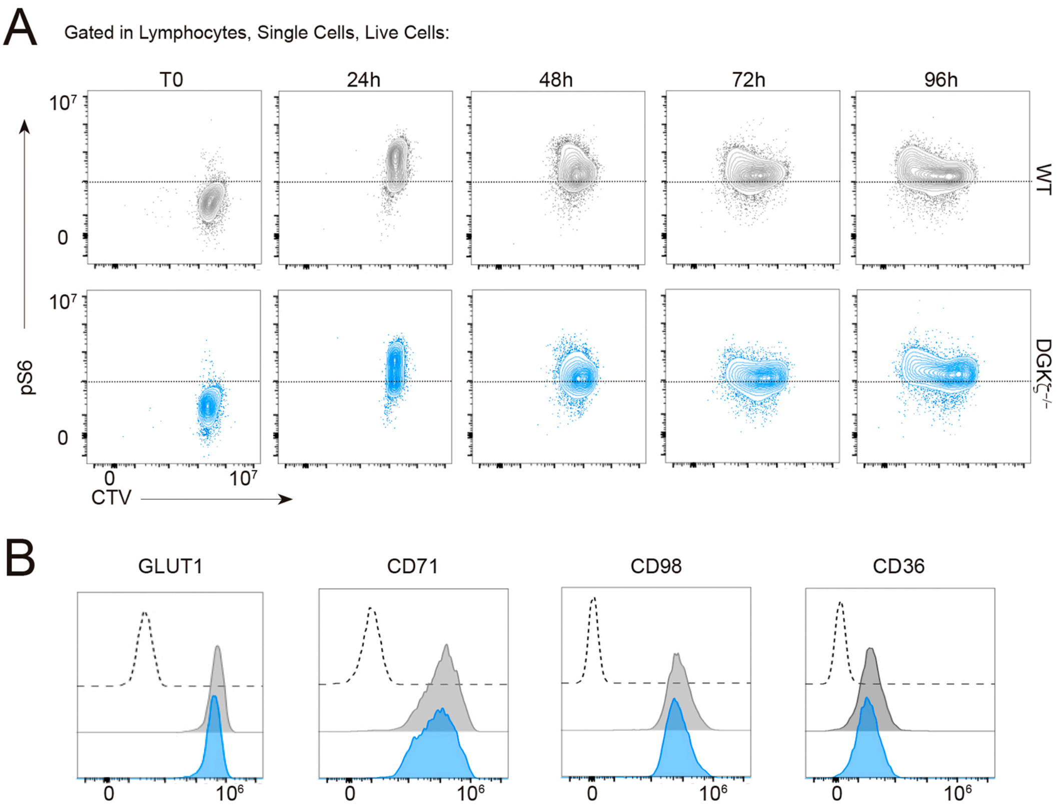


**Figure S3. DGKζ promotes anabolism and cell proliferation downstream CD40 signaling.** WT and DGKζ$^{-/-}$ (KO) B cells were cultured with anti-CD40/IL-4/IL-5, and analyzed at different time points by flow cytometry. **A,** Representative contour-plots of the phosphorylated form of S6 protein (p-S6) and CellTrace violet (CTV; to monitor cell proliferation) at each indicated time point; T0, freshly isolated cells. **B,** Expression profiles for the metabolite transporters GLUT1, CD71, CD98 and CD36 at 48 h in WT (grey) and DGKζ$^{-/-}$ (blue) B cells; dotted line, isotype control.

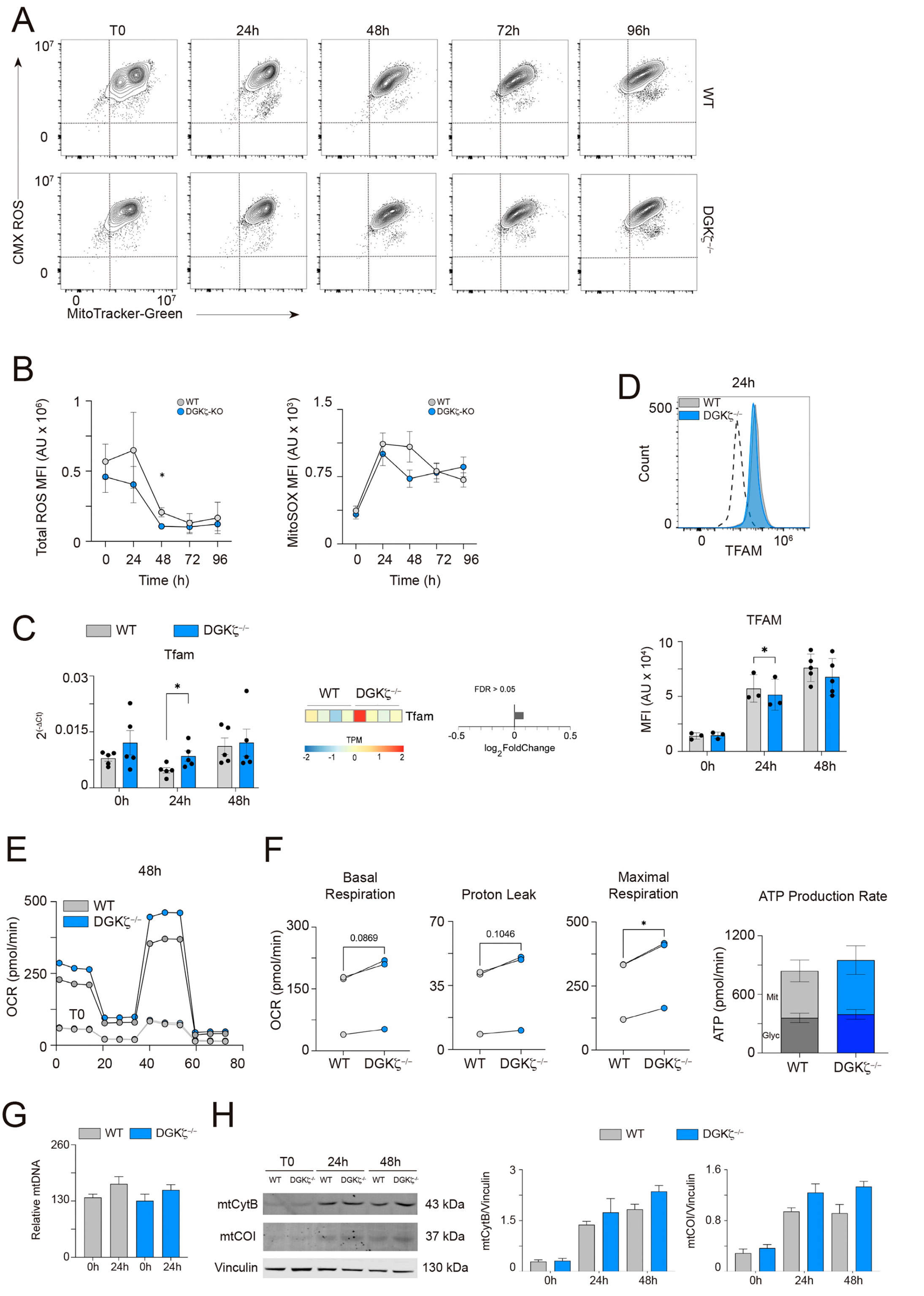


**Figure S4. DGKζ is essential for optimal mitochondrial function downstream of CD40 signaling in B cells.** Naïve and anti-CD40/IL-4/IL-5 stimulated WT and DGKζ$^{-/-}$ (KO) B cells were analyzed for parameters associated to mitochondrial function. **A**, Contour-plots to monitor mitochondrial mass (MitoTracker Green probe) and mitochondrial membrane potential (MitoTracker Red CMX-ROS probe) overtime. **B**, Total reactive oxygen species (ROS) production (CM-H2DCFDA probe), and mitochondrial ROS production (MitoSOX probe) in WT and DGK$\zeta^{-/-}$ B cells over time; mean fluorescence intensity (MFI) values in arbitrary units (AU) are shown. Data are the mean ±SEM of n=5 mice per genotype. **C**, mRNA levels of the *Tfam* gene measured by RT-qPCR in WT and DGK$\zeta^{-/-}$ B cells at the indicated time points; each dot corresponds to a mouse (left). Heatmap and bar-plot illustrating the expression profile (TPM, transcript per million mapped reads) and differential gene expression (log2FoldChange) of the *Tfam* gene obtained by RNA-seq; data from n=4 mice per genotype (right). **D,** Profile of TFAM protein expression detected in WT (grey) and DGK$\zeta^{-/-}$ (blue) B cells by intracellular staining at 24 h; dotted line, isotype control. MFI values for TFAM are shown in the bar-graph at the specified time points; each dot is a mouse. **E,** Profiles of oxygen consumption rate (OCR; in picomoles per minute, pmol/min) obtained by extracellular flux assays in WT and DGK$\zeta^{-/-}$ B cells at 48 h post-stimulation; profiles at time 0h (T0) are also shown. **F**, OCR values for different parameters related to mitochondrial respiration (basal and maximal respiration, proton leak) and values of ATP production rate from glycolytic (Glyc; dark color) and mitochondrial (Mit; light color) pathways; each dot corresponds to a mouse (n=3 per genotype). **G**, Mitochondrial DNA content (mtDNA) in WT and DGK$\zeta^{-/-}$ B cells at the indicated time points; data are the merge ±SEM of n=4 mice per genotype. **H**, Western blots for cytochrome B (CytB) and cytochrome oxidase subunit 1 (COI) mitochondrial proteins in B cells from both genotypes and at the indicated time points post-stimulation; vinculin, loading control. Quantification of the CytB and COI band intensities and normalized to that of vinculin; data are the mean ±SEM of n=3 mice per genotype. Statistical analysis: two-tailed paired parametric Student's *t*-test and, in C, edgeR for RNA-seq data. *, $p<0.05$.

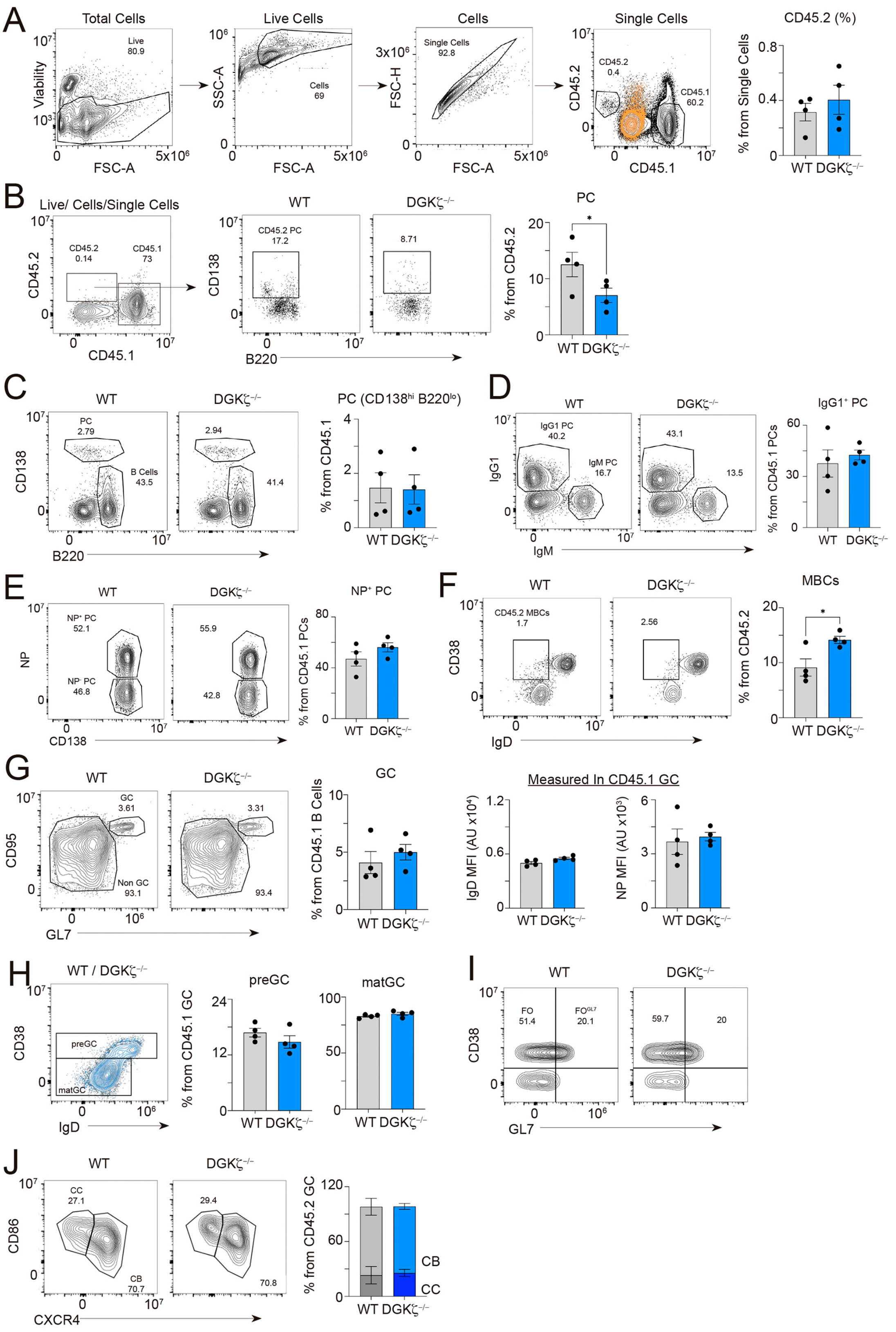


**Figure S5. DGKζ is needed for B cell progression in the germinal center response. A**, Gating strategy to analyze the populations generated at day 7 post-immunization in the spleen, in the CD45.1$^{+}$ host cells and transferred CD45.2$^{+}$ B cells. Orange overlayed plot, isotype control. Quantification of the $CD45.2^+$ cells frequencies detected at day 7 are shown in the graph. **B**, Bone marrow analysis of the $CD45.2^+$ plasma cell (PC) population. **C**, CD138/B220 contour-plots of CD45.1-gated cells in the spleen. The PC ($CD138^{hi}B220^{lo}$) and B cell populations are indicated. PC percentages are shown in the graph. **D,** IgG1/IgM contour-plots for $CD45.1^+$ PC; Graph, frequency values of $CD45.1^+$ $IgG1^+$ PC. **E,** NP/CD138 contour-plots for $CD45.1^+$ PC. $CD45.1^+$ $NP^+$ PC frequencies and mean fluorescence intensity (MFI) values for NP-antigen are shown in the graphs. **F**, CD38/IgD contour-plot of $CD45.2^+B220^+CD95^-GL7^-$ gated cells, showing the MBC population. Graph, frequency values of $CD38^+IgD^-$ MBCs. **G**, CD95/GL7 contour-plots of $CD45.1^+$ $B220^+$ gated B cells. The germinal center and non-GC B cell populations are shown. GC B cells frequency values, and MFI values for IgD and NP-antigen in the $CD45.1^+$ GC population are shown. **H**, Overlayed representative CD38/IgD contour-plots of $CD45.1^+$ GC B cells for WT (grey) and $DGK\zeta^{-/-}$ (blue) transferred mice. Frequency values of $CD45.1^+$ pre-GC and mature-GC B cells are shown. **I**, Gating strategy of $CD45.2^+$ follicular (FO) and $FO^{GL7}$ B cells. **J**, CD86/CXCR4 contour-plots of $CD45.2^+$ germinal center B cells, depicting the centroblast (CB) and centrocyte (CC) populations; frequency values shown on the graph. Statistical analysis: one sample t-test (C) and two-tailed unpaired (D, E) or paired (G, H) parametric Student's *t*-test. Data shown in bar graphs include the mean value ± SEM for each case; each dot in all the graphs corresponds to a mouse.

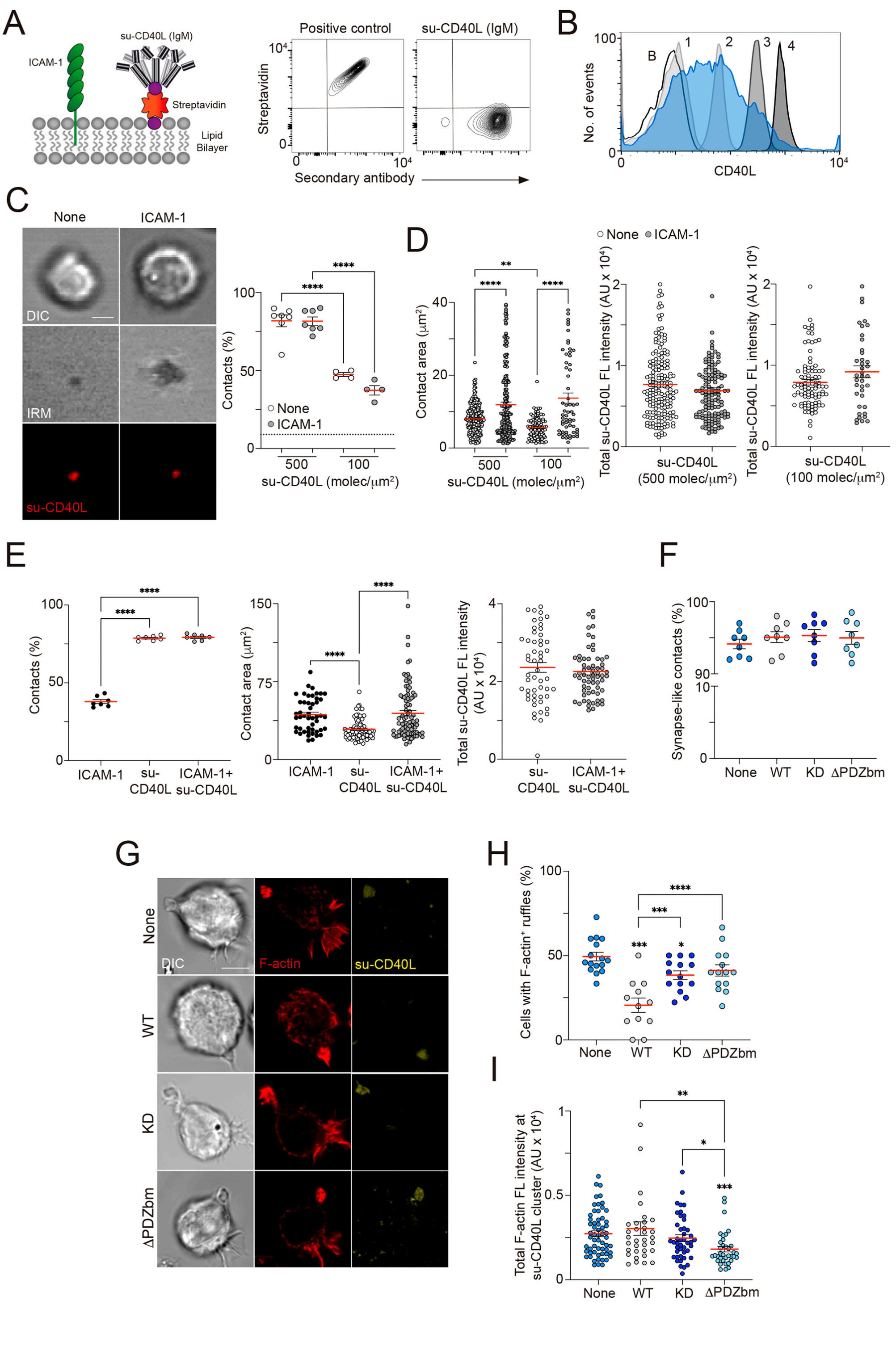


**Figure S6. DGKζ drives F-actin polymerization and LFA-1-mediated adhesion at the CD40-mediated synapse-like contact.** **A**, Scheme of the artificial planar lipid bilayer containing GPI-linked ICAM-1 and biotinylated lipids, the latter are used to tether fluorescently labeled streptavidin followed by mono-biotinylated anti-mouse CD40 (su-CD40L) (left). Countor-plots of fluorescently-labeled streptavidin and goat anti-hamster secondary antibody for streptavidin-beads loaded with a poly-biotinylated antibody (positive control) and the mono-biotinylated anti-mouse CD40 (su-CD40L) (right). **B**, Profile of standard beads with a defined rat IgG-binding capacity (B, none; 1 to 4, see methods; grey-filled histograms) and overlayed in blue the CD40L profile of mouse blast T cells, to estimate the molecular density (molecules/$\mu m^2$) of CD40L at the activated T cell surface **C-D**, WT B cells were settled on bilayers containing su-CD40L and in absence or presence of ICAM-1 for 30 min before being imaged. **C**, DIC, IRM and fluorescence su-CD40L images at the contact plane. Percentage of B cells with established synapse-like contact (estimated by IRM and fluorescence) are shown in the graph; dotted line, basal adhesion frequency on ICAM-1 containing bilayers. **D**, Values of the area of contact (estimated by IRM) and of the total su-CD40L fluorescence (FL) intensity accumulated at the synapse-like contact (in arbitrary units, AU) at the specified su-CD40L densities (each density was imaged using different acquisition conditions of laser gain). **E**, A20 cells settled on bilayers as in C (su-CD40L 500 molecules/$\mu m^2$). Frequency of contacts, contact area and total su-CD40L FL intensity values at the synapse-like contact are shown for each indicated condition. **F-H**, A20 cells transfected with the GFP-fused DGKζ constructs (WT; KD; ΔPDZbm) on lipid bilayers containing ICAM-1 and su-CD40L (500 molecules/$\mu m^2$) for 30 min, and then were imaged or fixed for immunofluorescence. **F**, Percentages of cells with synapse-like contact; GFP negative transfected cells were used as controls (None). **G**, DIC, and fluorescence F-actin and su-CD40L images of A20 cells. **H**, Percentages of A20 cells showing F-actin-enriched membrane ruffles. **I**, Quantification of the total F-actin FL intensity accumulated at the su-CD40L cluster. Each dot in C, E (left graph), F, and H corresponds to an imaged field, and to a single cell in the rest. Data shown in C, E corresponds to a representative experiment; data shown in D, F, H, and I are the mean ±SEM of n=3 experiments. Statistical analysis: two-tailed unpaired Student's *t*-test (D, mid- and right graph; E, right graph) and One-way ANOVA (rest of the data). *, $p<0.05$; **, $p<0.01$; ***, $p<0.001$; ****, $p<0.0001$.

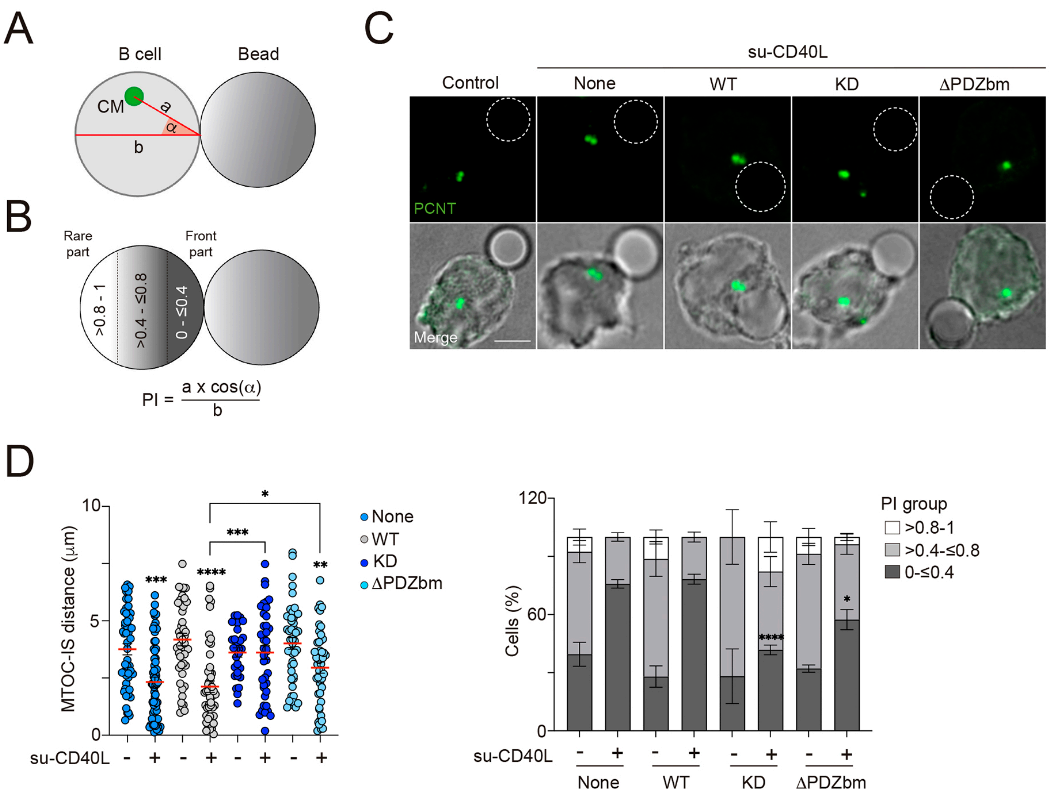


**Figure S7. Analysis of organelle translocation to the CD40-driven synapse-like contact. A**, Scheme for the calculation of the distance of organelles (MTOC, lysosomes, mitochondria) to the synapse contact (see methods). **B**, Estimation of the Polarity Index (PI) value and illustration of the three groups established based on the PI value. **C**, Fluorescence pericentrin and merged DIC/pericentrin images of A20 cells transfected with the DGKζ constructs (WT; KD; ΔPDZbm) forming conjugates with su-CD40L loaded beads or unloaded (control); None, GFP negative cells. **D**, Left, Distance values of the MTOC (identified by pericentrin staining) to the synapse contact per cell, calculated as depicted in A, for each transfectant. Right, frequency of A20 B cells distributed in each PI group. Data shown are the mean ±SEM of n=3 experiments. Statistical analysis: One-way (D, left graph) and two-way ANOVA (D, right graph). *, $p$<0.05; **, $p$<0.01; ***, $p$<0.001; ****, $p$<0.0001.

### SUPPLEMENTAL MOVIES

**Movie S1. Plasma membrane ruffling in B cells with established CD40-mediated synapse-like contact**. Movies of representative WT (left) and DGKζ$^{-/-}$ (right) B cells with a CD40-driven synapse formed on the lipid bilayer system. B cells were incubated with artificial membranes containing GPI-linked ICAM-1 and su-CD40L (500 molecules/μm$^2$) for 30 min, at 37ºC, and then imaged to monitored cell behavior (movie length, 5 min; images acquired every 15 seconds). DIC and IRM images are shown.

**Movie S2. Cell behavior of A20 B cells expressing DGKζ constructs.** Movies of representative A20 B cells expressing the GFP-tagged DGKζ-WT, -KD or -ΔPDZbm construct, or GFP negative (None), with a CD40-driven synapse formed on the lipid bilayer system. A20 B cells were incubated with artificial membranes containing GPI-linked ICAM-1 and su-CD40L (500 molecules/μm$^2$) for 20 min, at 37ºC, and then imaged to monitored cell behavior (movie length, 5 min; images acquired every 15 seconds). DIC and IRM images are shown.

### SUPPLEMENTAL METHODS

#### RNA-seq Data Bioinformatic Analysis

For analysis of the RNA-seq data, paired-end reads were mapped to a mouse reference genome (GRCm38/mm10 assembly) using BWA-MEM 0.7.15 (http://biobwa.sourceforge.net). Alignments were formatted to BAM and duplicates were removed with Picard tools 2.9.0 (http://broadinstitute.github.io/picard/). Relative expression of transcripts was quantified with StringTie 1.3.3[1], converted to transcripts per million (TPM) reads, and kept for later analysis when TPM > 0 in all samples. The expression level of all genes was estimated by edgeR[2]. Genes with a Benjamini–Hochberg adjusted P-value <0.05 and a cut-off of 0.6 in log2(fold change) were considered differentially expressed (DEG). RNA-Seq data are deposited in the Gene Expression Omnibus (GEO) under accession code GSE283915. For bioinformatics and Functional Enrichment, data were obtained, processed and annotated using R (R Development Core Team, 2014) and Bioconductor programs[3]. Functional enrichment analysis was performed using GSEA (https://www.gsea-msigdb.org/)[4], and heatmaps were built using Pheatmap.

REFERENCES

1. Pertea, M., Pertea, G.M., Antonescu, C.M., Chang, T.C., Mendell, J.T., and Salzberg, S.L. (2015). StringTie enables improved reconstruction of a transcriptome from RNA-seq reads. Nat Biotechnol *33*, 290-295. 10.1038/nbt.3122.
2. Robinson, M.D., McCarthy, D.J., and Smyth, G.K. (2010). edgeR: a Bioconductor package for differential expression analysis of digital gene expression data. Bioinformatics *26*, 139-140. 10.1093/bioinformatics/btp616.
3. Huber, W., Carey, V.J., Gentleman, R., Anders, S., Carlson, M., Carvalho, B.S., Bravo, H.C., Davis, S., Gatto, L., Girke, T., et al. (2015). Orchestrating high-throughput genomic analysis with Bioconductor. Nat Methods *12*, 115-121. 10.1038/nmeth.3252.
4. Subramanian, A., Tamayo, P., Mootha, V.K., Mukherjee, S., Ebert, B.L., Gillette, M.A., Paulovich, A., Pomeroy, S.L., Golub, T.R., Lander, E.S., and Mesirov, J.P. (2005). Gene set enrichment analysis: a knowledge-based approach for interpreting genome-wide expression profiles. Proc Natl Acad Sci U S A *102*, 15545-15550. 10.1073/pnas.0506580102.
